# A STAG2-PAXIP1/PAGR1 axis suppresses lung tumorigenesis

**DOI:** 10.1101/2024.09.14.613043

**Authors:** Emily L. Ashkin, Yuning J. Tang, Haiqing Xu, King L. Hung, Julia Belk, Hongchen Cai, Steven Lopez, Deniz Nesli Dolcen, Jess D. Hebert, Rui Li, Paloma A. Ruiz, Tula Keal, Laura Andrejka, Howard Y. Chang, Dmitri A. Petrov, Jesse R. Dixon, Zhichao Xu, Monte M. Winslow

**Affiliations:** Cancer Biology Program, Stanford University School of Medicine, Stanford, CA, USA 94305; Department of Genetics, Stanford University School of Medicine, Stanford, CA, USA 94305; Department of Biology, Stanford University, Stanford, CA, USA 94305; Gene Expression Laboratory; Salk Institute for Biological Studies, La Jolla, CA, USA 92037; Center for Personal Dynamic Regulomes, Stanford University School of Medicine, Stanford, CA, USA 94305; Howard Hughes Medical Institute, Stanford University School of Medicine, Stanford, CA, USA 94305; Chan Zuckerberg Biohub, San Francisco, CA, USA 94158; Department of Pathology, Stanford University School of Medicine, Stanford, CA, USA 94305

## Abstract

The cohesin complex is a critical regulator of gene expression. *STAG2* is the most frequently mutated cohesin subunit across several cancer types and is a key tumor suppressor in lung cancer. Here, we coupled somatic CRISPR-Cas9 genome editing and tumor barcoding with an autochthonous oncogenic KRAS-driven lung cancer model and show that STAG2 is uniquely tumor suppressive among all core and auxiliary cohesin components. The heterodimeric complex components *PAXIP1* and *PAGR1* have highly correlated effects with *STAG2* in human lung cancer cell lines, are tumor suppressors *in vivo*, and are epistatic to *STAG2* in oncogenic KRAS-driven lung tumorigenesis *in vivo*. STAG2 inactivation elicits changes in gene expression, chromatin accessibility and 3D genome conformation that impact cancer cell state. Gene expression and chromatin accessibility similarities between STAG2- and PAXIP1-deficient neoplastic cells further relates STAG2-cohesin to PAXIP1/PAGR1. These findings reveal a STAG2-PAXIP1/PAGR1 tumor-suppressive axis and uncover novel PAXIP1-dependent and PAXIP1-independent STAG2-cohesin mediated mechanisms of lung tumor suppression.

**SUMMARY:** STAG2 is a frequently mutated cohesin subunit across several cancers and one of the most important functional suppressors of lung adenocarcinoma. Our findings underscore important roles of STAG2 in suppressing lung tumorigenesis and highlight a STAG2-PAXIP1/PAGR1 tumor-suppressive program that may transcend cancer type.

## INTRODUCTION

The cohesin complex is a large multi-subunit complex that plays important roles in regulating genome conformation, gene expression and determining cell fate (1). There are two classes of cohesin complexes, distinguished by whether they contain the paralogs STAG1 or STAG2 (2). STAG1- and STAG2-cohesin are thought to control different gene expression programs (3–5). STAG1-cohesin colocalizes with CTCF to form topologically associating domain boundaries that drive long-range chromatin interactions for gene transcription (5,6). Conversely, STAG2-cohesin has been shown to drive mid-range chromatin interactions (5,7); however, it remains unknown whether additional proteins contribute specifically to STAG2-cohesin regulated gene expression (8,9).

Notably, *STAG2* is the most frequently mutated cohesin complex member across many cancer types, including bladder cancer, Ewing’s sarcoma, acute myeloid leukemia (AML), and lung adenocarcinoma (9–18). Using *in vivo* CRISPR/Cas9 screens within genetically engineered mouse models of lung cancer, we and others found STAG2 to be an important tumor suppressor in lung adenocarcinoma (19,20). *STAG2* mutations occur in ∼4% of human lung adenocarcinomas, and STAG2 protein expression is low or absent in ∼20% of these tumors (19). In lung cancer, STAG2-mediated tumor suppression is not through an impact on chromosomal instability or hindering MAPK signaling, as previously suggested for other cancer types (10,19,21–25). Thus, the mechanisms through which *STAG2* inactivation drives lung tumor growth remain largely unknown and crucial for understanding cellular processes that contribute to tumor suppression.

Here, using CRISPR/Cas9-mediated somatic genome editing within genetically engineered mouse models of lung adenocarcinoma, we investigate the tumor-suppressive capacity of STAG2, every other component of the cohesin complex, and potential co-operators of STAG2-cohesin in lung cancer. We identify PAXIP1 and PAGR1, which form an obligatory heterodimer that has been suggested to be involved in DNA damage repair, cell cycle control, and transcription regulation (26–29) as potent lung tumor suppressors, and provide multiple complementary lines of evidence for a STAG2-PAXIP1/PAGR1 tumor-suppressive axis.

## RESULTS

### Lung tumor suppression by STAG2 is unique among cohesin subunits

STAG2 is a tumor suppressor in oncogenic KRAS-driven lung cancer, and other cohesin subunits are frequently mutated in KRAS-driven lung cancer as well as in several other cancer types (9–14,19,30–33). To determine whether the inactivation of other cohesin subunits would also increase autochthonous lung tumor growth, we used tumor barcoding and somatic CRISPR/Cas9-mediated genome editing (19,34). We used a recently optimized version of tumor barcoding coupled with high-throughput barcode sequencing (Tuba-seq), in which lentiviral vectors have a diverse barcode (BC) integrated within the U6 promoter directly 5’ of the sgRNA (Tuba-seq^Ultra^; U6 barcode Labeling with per-Tumor Resolution Analysis; **Fig. 1a-b** and **Methods**) (35). Tuba-seq^Ultra^ enables the quantification of the size of each clonal tumor and the number of tumors with each sgRNA based on amplification of the BC-sgRNA region from bulk tumor-bearing lungs followed by high-throughput sequencing.

**Figure 1.**
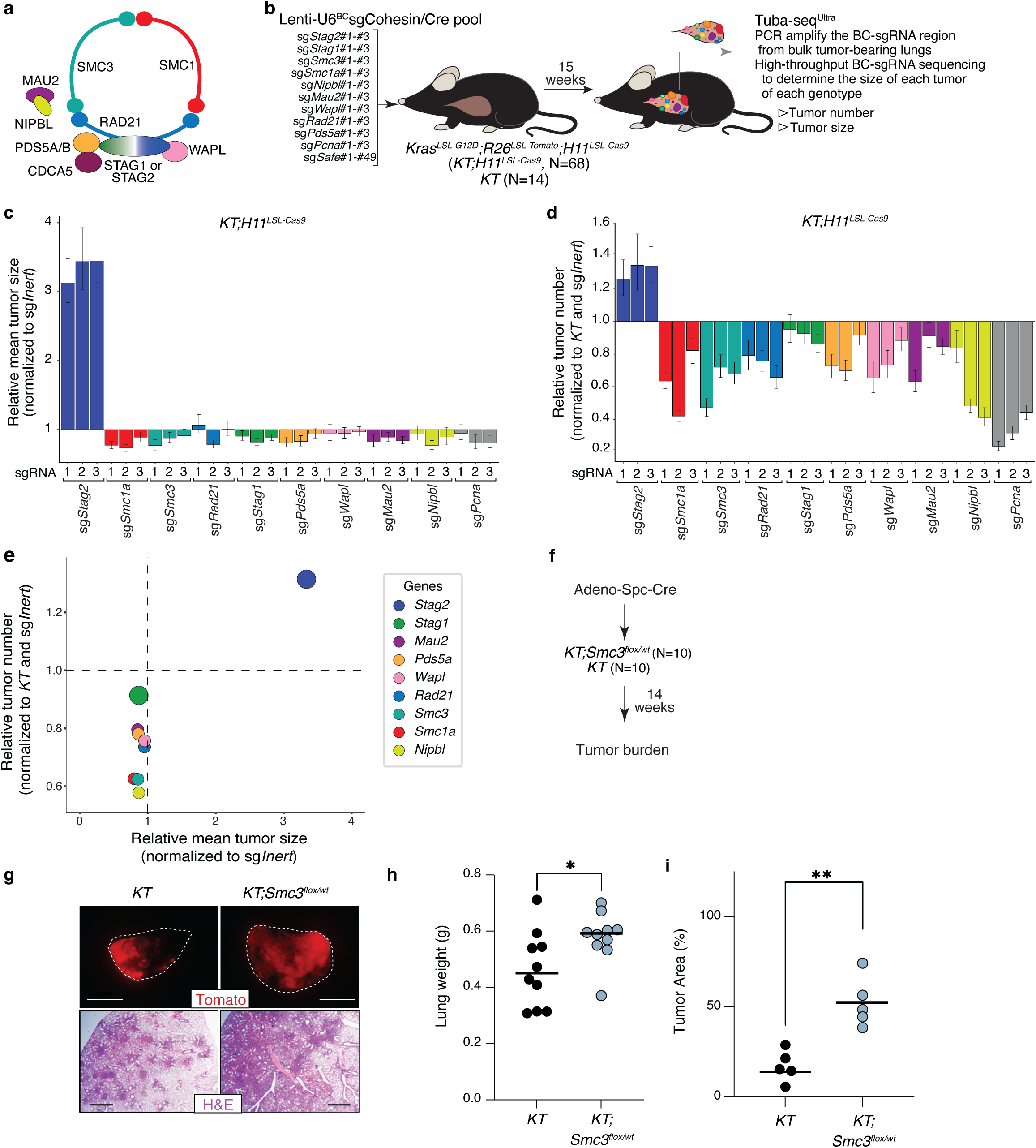
Inactivation of *Stag2* and heterozygous inactivation of the cohesin suabunit *Smc3* uniquely increase lung tumor growth. **a)** Schematic of the cohesin complex with subunits labeled. **b)** Tumor initiation with a pool of Lenti-U6^BC^sgRNA/Cre vectors. Genotype and number of mice are indicated. Tuba-seq^Ultra^ was performed on each tumor-bearing lung 15 weeks after tumor initiation, followed by analyses to quantify tumorigenesis. **c)** Relative mean tumor size (normalized to sg*Inert*). Mean +/− 95% confidence intervals are shown. Dotted line indicates no effect. **d)** Relative tumor number (normalized to *KT* and sg*Inert*). Mean +/− 95% confidence intervals are shown. Dotted line indicates no effect. **e)** Relative tumor number (normalized to *KT* and sg*Inert*) correlated with relative mean tumor size (normalized to sg*Inert*). Data represents one replicate of two independent experiments. **f)** Tumor initiation with Adeno-Spc-Cre in *KT* and *KT;Smc3^flox/+^* mice. Mouse number is indicated. **g)** Representative fluorescence and histology images of lung lobes from the indicated genotypes of mice. Top scale bars, 5 mm, and bottom scale bars, 1 mm. h) Lung weights from *KT* and *KT;Smc3^flox/+^* mice. Each dot represents a mouse and the bar is the mean. Data are representative of two independent experiments. **i)** Percentage of Tomato-positive tumor area detected via histology. Each dot represents a mouse and the bar is the mean. *p-value < 0.05, ** p-value < 0.01, via unpaired t-test. Raw values and significance of each effect is shown in Supplementary Table 1.

We generated a lentiviral pool that contained barcoded lentiviral-sgRNA/Cre Tuba-seq^Ultra^ vectors targeting each cohesin gene, an essential gene, and Safe-cutting “inert” sgRNAs (Lenti-U6^BC^sgCohesin/Cre with 3 sgRNAs/gene; **Fig. 1b, Methods** and **Supplementary Table S1**). We initiated tumors with Lenti-U6^BC^sgCohesin/Cre in *Kras^LSL-G12D/+^;R26^LSL-Tomato^;H11^LSL-Cas9^* (*KT;H11^LSL-Cas9^*) mice and Cas9-negative *Kras^LSL-G12D/+^;R26^LSL-Tomato^* (*KT*) mice (**Methods**). After fifteen weeks of tumor growth, we extracted DNA from bulk tumor-bearing lungs, PCR amplified the BC-sgRNA region, and high-throughput sequenced the amplicon. By tallying the number of reads from each BC-sgRNA and normalizing to “spike-in” control cells with a known BC-sgRNA that were added at a defined cell number, we quantified the number of neoplastic cells in each clonal tumor with each sgRNA (**Methods**).

We calculated tumor number (the number of BCs associated with each sgRNA normalized to the expected number from Cas9-negative *KT* mice) and several metrics of tumor size (log-normal mean tumor size and tumor sizes at defined percentiles within the tumor size distribution; **Methods**). Consistent with previous observations, *Stag2* inactivation greatly increased tumor size and modestly increased tumor number (**Fig. 1c-d**) (19,20). Inactivation of most of the other cohesin complex components greatly reduced tumor number and tumor size, although not to the same extent as inactivation of the known essential gene *Pcna* (**Fig. 1c-e**). All three sgRNAs targeting *Stag1* only modestly reduced tumor number and tumor size. Collectively, these data are consistent with cohesin in general having an essential function and the well-described ability of STAG2 can compensate for STAG1 in this essential function (3,6,21,36) (**Fig. 1c-e**). Importantly, these results provided no indication that STAG1 is a tumor suppressor and are consistent with distinct roles of STAG1- and STAG2-cohesin in lung cancer.

### Heterozygous inactivation of the cohesin subunit *Smc3* increases lung tumorigenesis

While homozygous inactivation of cohesin subunits is detrimental to lung tumorigenesis, mutations in these genes are common in lung cancer, and heterozygous inactivation of core cohesin genes increase tumorigenesis in other contexts (37,38). Thus, we hypothesized that heterozygous inactivation of core cohesin subunits could increase lung tumor growth. We chose to focus on the core cohesion component Smc3 due to the availability of mice with an *Smc3* floxed allele (39). To generate tumors with inactivation on one allele of *Smc3*, we initiated lung tumors in *KT;Smc3^flox/+^*and *KT* control mice. After fourteen weeks of tumor growth, *KT;Smc3^flox/+^* mice had significantly greater tumor burden and higher tumor number compared to *KT* mice as assessed by total lung weight, fluorescent imaging, and histology **(Fig. 1f-i)**. These genetic data support a model in which the net effect of a reduction in cohesin is increased lung tumor growth potentially driven by trade-offs between a reduction in STAG2 cohesin-mediated tumor suppression and the essential function of cohesin in general.

### DepMap and *in vivo* analyses reveal cooperation between *STAG2* and the PAXIP1/PAGR1 complex in KRAS-driven lung tumors

To understand mechanistically how STAG2-cohesin regulates tumorigenesis, we next sought to find genes that cooperate with STAG2 to mediate these effects. We analyzed the CRISPR/Cas9 human cancer cell line Dependency Map data for genes whose effects of cell growth (referred to as their gene effect) were most similar to that of *STAG2* (**Methods**) (40,41). We initially assessed the impact of inactivating each gene across all cell lines (excluding those with *STAG2* mutations, N=1018) and calculated the correlation (Pearson’s r) with the *STAG2* gene effect. Consistent with previous analyses, *PAXIP1* and its obligate heterodimer partner *PAGR1* had the most highly correlated gene effects with *STAG2* (r=0.62 and 0.61, respectively) (8,42) and their correlation with *STAG2* was ∼2-fold higher than any other gene, including core and auxiliary cohesin subunits (**Fig. 2a** and **Supplementary Fig. S1a-c**). Importantly, in oncogenic KRAS^G12/G13^-driven cell lines (N=120), as well as in oncogenic KRAS^G12/G13^-driven lung adenocarcinoma cell lines (N=22), the gene effects of *PAXIP1* and *PAGR1* robustly correlated with the gene effect of *STAG2* (**Fig. 2b-c** and **Supplementary Fig. S1d-e**). Finally, the positive correlations between *STAG2* and *PAXIP1/PAGR1* gene effects extended to other cancer types where *STAG2* is frequently mutated, including acute myeloid leukemia, bladder cancer, and Ewing’s sarcoma (**Supplementary Fig. S3f-i**) (9–14).

**Figure 2.**
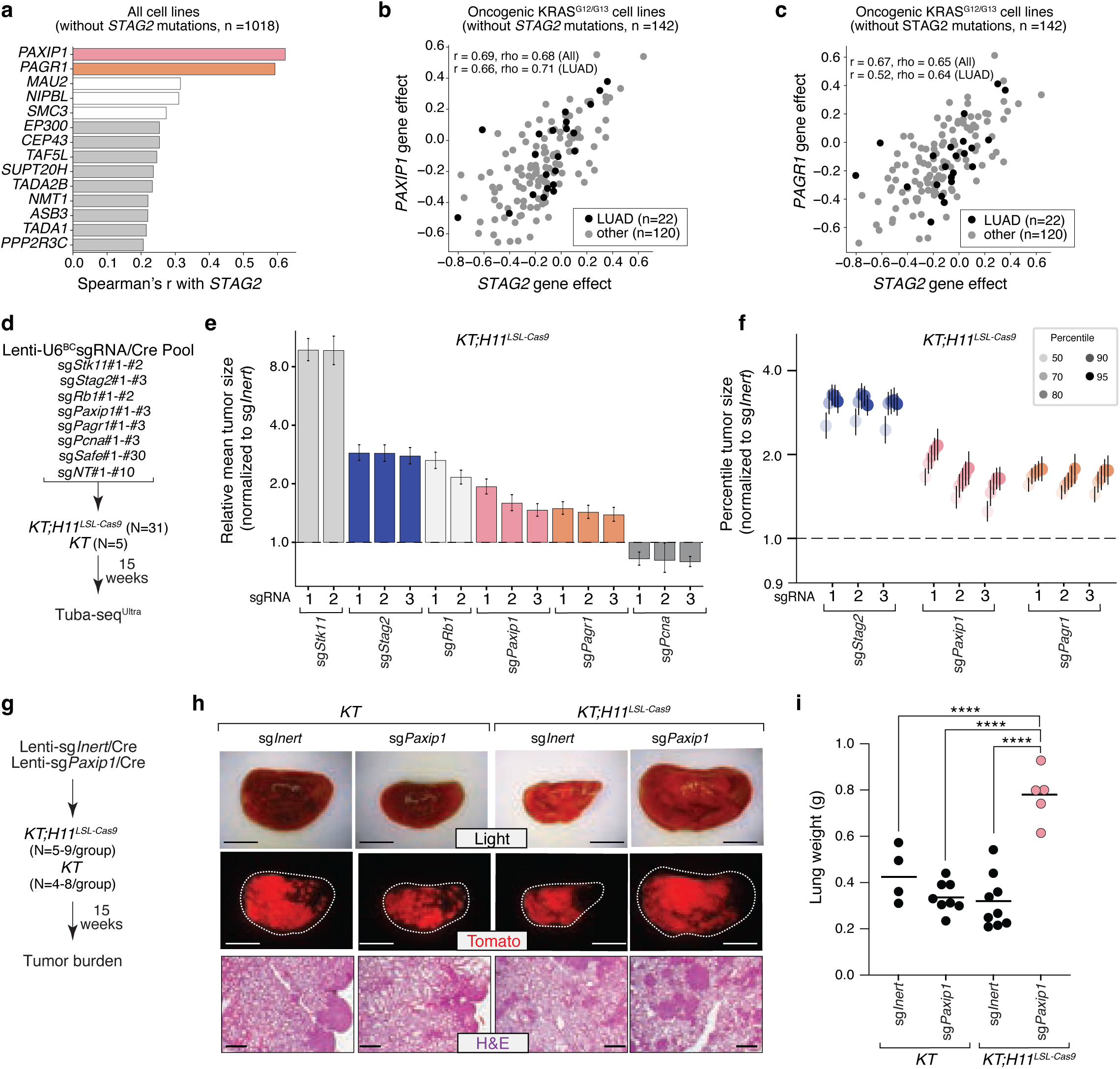
STAG2, PAXIP1, and PAGR1 effects are highly correlated in human cancer cell lines and PAXIP1/PAGR1 are tumor suppressive in KRAS-driven lung tumors *in vivo*. **a)** Genes with the most correlated effects with *STAG2* inactivation from the Dependency Map (DepMap). Cell lines with *STAG2* mutations were excluded. *PAXIP1* and *PAGR1* are colored bars. Core cohesin complex genes are white bars. **b-c)** Gene effects for *PAXIP1* and *STAG2* inactivation **(b)** and for *PAGR1* and *STAG2* inactivation **(c)**. Each dot represents a cell line with an oncogenic mutation at codons 12 or 13 of KRAS. Cell lines with *STAG2* mutations were excluded. Lung adenocarcinoma (LUAD) cell lines are shown as black dots. Spearman’s r and Pearson rho for all cell lines and for LUAD cell lines are indicated. **d)** Tumor initiation with a pool of barcoded Lenti-U6^BC^sgRNA/Cre Tuba-seq^Ultra^ vectors. Genotype and number of mice are indicated. Tuba-seq^Ultra^ was performed on each tumor-bearing lung 15 weeks after tumor initiation, followed by analyses to quantify tumorigenesis. **e)** Relative mean tumor size (normalized to sg*Inert*). Mean +/− 95% confidence intervals are shown. Dotted line indicates no effect. **f)** Tumor sizes at the indicated percentiles for tumors with sgRNA targeting *Stag2*, *Paxip1*, or *Pagr1* (normalized to sg*Inert*) in *KT;H11^LSL-Cas9^* mice. Each gene was targeted with three sgRNAs. Error bars indicate 95% confidence intervals. Dotted line indicates no effect. Data represents one replicate of two independent experiments. **g)** Tumor initiation with the indicated Lenti-sgRNA/Cre vectors in separate cohorts of *KT* and *KT;H11^LSL-Cas9^* mice. Number of mice in each group is indicated. Tumor burden was quantified 15 weeks after tumor initiation. **h)** Representative brightfield and fluorescence images of lung lobes and histology from the indicated groups of mice. Top scale bars and middle scale bars, 5 mm. Lower scale bars, 500 μm. **i)** Lung weights of mice in each group. Each dot represents a mouse and the bar is the mean. ****p-value <0.0001 by unpaired t-test. Data represents one replicate of two independent experiments. Raw values and significance of each effect is shown in Supplementary Table 2.

To determine whether PAXIP1 and PAGR1 are tumor suppressors in lung cancer, we used Tuba-seq^Ultra^ to quantify the impact of inactivating *STAG2, PAXIP1,* or *PAGR1* on oncogenic KRAS-driven lung tumorigenesis (**Fig. 2d**). We initiated tumors *KT;H11^LSL-Cas9^* and *KT* mice with a pool of Lenti-U6^BC^-sgRNA/Cre vectors that contained sgRNAs targeting each gene of interest (**Fig. 2d** and **Supplementary Table S2**). After fifteen weeks of tumor growth, we quantified the number of neoplastic cells in each tumor using Tuba-seq^Ultra^ (**Fig. 2d**). Inactivation of *Paxip1* and *Pagr1* significantly increased tumor size, on par with the effect of inactivating the canonical tumor suppressor *Rb1* (**Fig. 2e**). Inactivation of *Paxip1* and *Pagr1* showed modest effects on tumor number, on par with inactivation of *Stk11*, *Stag2*, and *Rb1* (**Supplementary Fig. S1j)**. The tumor-suppressive effects of PAXIP1 and PAGR1 were less than that of STAG2 (**Fig. 2e-f**). No vectors impacted tumorigenesis in the absence of Cas9 (**Supplementary Table S2**).

To validate the tumor-suppressive effects of PAXIP1, we initiated lung tumors with Lenti-sg*Inert*/Cre and Lenti-sg*Paxip1*/Cre in separate cohorts of *KT;H11^LSL-Cas9^* and *KT* mice (**Fig. 2g**). PAXIP1 inactivation was confirmed by western blotting on sorted neoplastic cells (**Supplementary Fig. S1k**). Overall tumor burden as assessed by lung weight was significantly higher in *KT;H11^LSL-Cas9^* sg*Paxip1* mice compared to each of the three control cohorts (mice with *KT;H11^LSL-Cas9^* sg*Inert*, *KT* sg*Paxip1*, and *KT* sg*Inert* tumors)**(Fig. 2h-i)**. Collectively, these quantitative *in vivo* data and DepMap findings from human cancer cell lines show that STAG2-cohesin effects correlate with PAXIP1/PAGR1 effects in lung tumorigenesis (**Fig. 2a-f**).

### STAG2 inactivation modifies the overall tumor suppressive landscape and is epistatic to PAXIP1 and PAGR1

To determine whether PAXIP1 and PAGR1 function in the same pathway as STAG2, as well as extend our understanding of how STAG2-deficiency changes the impact of coincident genetic alterations on KRAS-driven lung tumorigenesis, we quantify the impact of inactivating a curated list of genes on tumor initiation and growth in *Stag2*-proficient and *Stag2-*deficient tumors using Tuba-seq^Ultra^ **(Fig. 3a-b)**. We generated a pool of Lenti-U6^BC^-sgRNA/Cre vectors that targeted ∼160 genes, including those that correlate with *STAG2* effects in DepMap (e.g. *Paxip1* and *Pagr1*), several canonical tumor suppressor genes, genes with protein-protein interactions with STAG2, and genes downstream of STAG2 based on preliminary molecular analyses (**Fig. 3a-b**, **Supplementary Table S3,** and **Methods)**. Each gene was targeted with 3-6 sgRNAs, and this pool contained control Safe-cutting “inert” sgRNAs and sgRNAs targeting several control essential genes (**Fig. 3a,b**). We initiated lung tumors in *KT, KT;H11^LSL-Cas9^* and *KT;H11^LSL-Cas9^;Stag2^flox^* mice (**Fig. 3b**). After thirteen weeks of tumor growth, we used Tuba-seq^Ultra^ to quantify the number of neoplastic cells in each tumor with each sgRNA (**Fig. 3b**).

**Figure 3.**
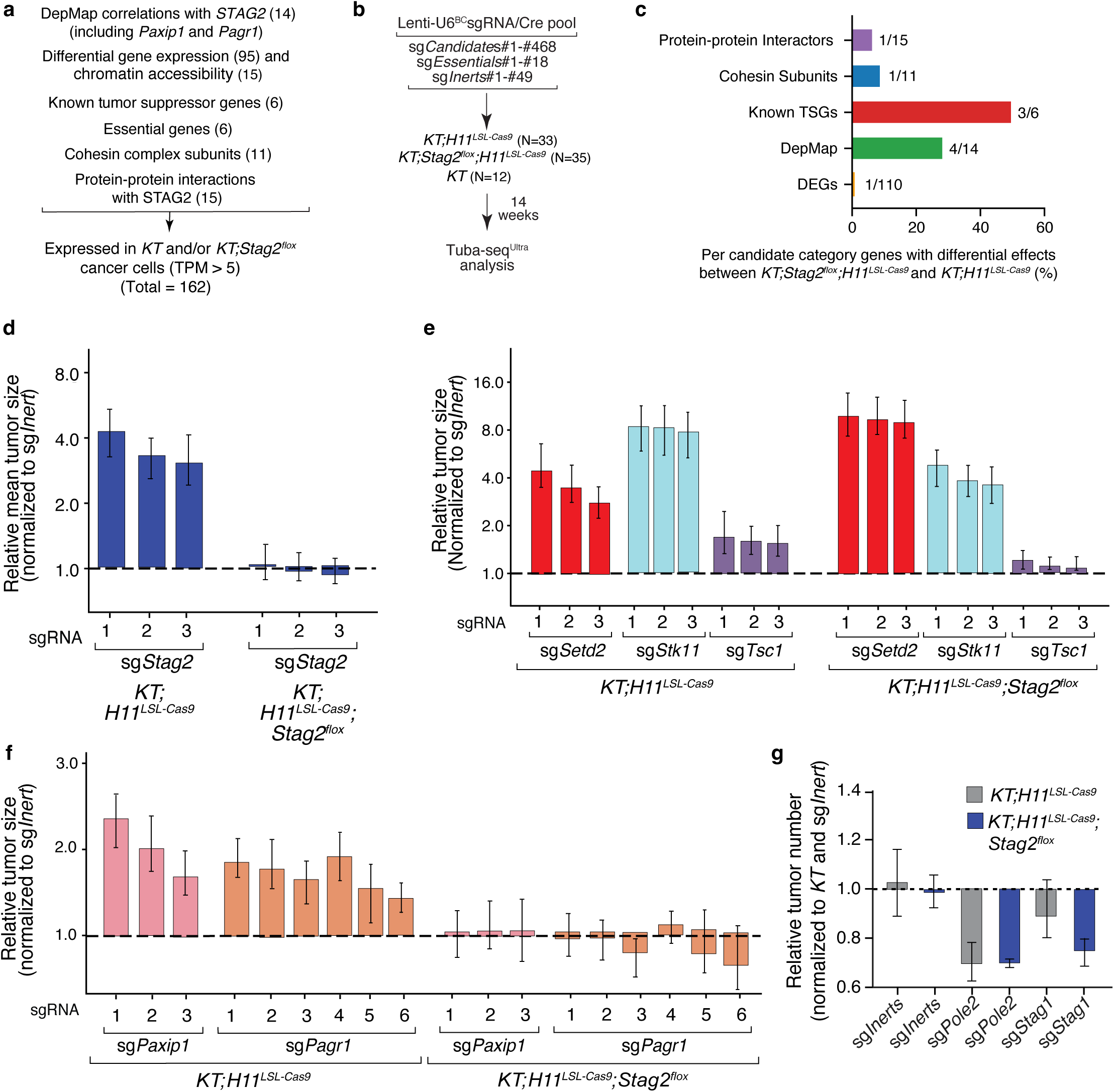
Genetic interactions with STAG2 include modification of overall tumor suppression and functional dependency with PAXIP1 and PAGR1. **a)** List of candidate gene criteria for pool of 468 barcoded Lenti-sgRNA/Cre vectors. Tumor initiation with Lenti-U6^BC^-sgRNA/Cre vectors in *KT;H11^LSL-Cas9^, KT;^H11LSL-Cas9^;Stag2^flox^*, and *KT* mice. Number of mice in each group is indicated. Sequencing was performed on each tumor-bearing lung 14 weeks after tumor initiation, followed by analysis to quantify tumorigenesis. **b)** Bar plot with percent of genes in each candidate category with differential effects between *KT;H11^LSL-Cas9^* and *KT;H11^LSL-Cas9^;Stag2^flox^*. **c)** Relative mean tumor size of tumors with sgRNAs targeting *Stag2* (normalized to sg*Inert*) in *KT;H11^LSL-Cas9^* mice compared to *KT;H11^LSL-Cas9^;Stag2^flox^*mice. Mean +/− 95% confidence intervals are shown. Dotted line indicates no effect. **d)** Relative mean tumor size of tumors with sgRNAs targeting *Setd2*, *Stk11*, or *Tsc1* (normalized to sg*Inert*) in *KT;H11^LSL-Cas9^* mice compared to *KT;H11^LSL-Cas9^;Stag2^flox^*mice. Mean +/− 95% confidence intervals are shown. Dotted line indicates no effect. **e)** Relative mean tumor size of tumors with sgRNAs targeting *Paxip1* or *Pagr1* (normalized to sg*Inert*) in *KT;H11^LSL-Cas9^* mice compared to *KT;H11^LSL-Cas9^;Stag2^flox^*mice. Mean +/− 95% confidence intervals are shown. Dotted line indicates no effect. **g)** Comparison of average relative tumor number for sgRNAs targeting *Stag1*, *Stag2*, *Pole2* (essential gene), or *Inerts* in *KT;H11^LSL-Cas9^* mice and *KT;H11^LSL-Cas9^;Stag2^flox^* mice. Mean +/− 95% confidence intervals are shown. Raw values and significance of each effect is shown in Supplementary Table 4. Data for *KT;H11^LSL-Cas9^* and *KT* mice represent one replicate of two independent experiments.

*Stag2* deficiency in *KT;H11^LSL-Cas9^;Stag2^flox^*mice significantly changes the impact of inactivating only a small number of genes, including several canonical tumor suppressor genes and genes identified through DepMap (**Fig. 3c**). As expected, sgRNAs targeting *Stag2* increased tumor growth *KT;H11^LSL-Cas9^* mice but not in *KT;H11^LSL-Cas9^;Stag2^flox^* mice (**Fig. 3d**). Inactivation of *Setd2* increased the growth of *Stag2*-deficient tumors more than *Stag2*-proficient tumors, while inactivation of *Stk11* or *Tsc1* increased the growth of *Stag2*-deficient tumors less than *Stag2*-proficient tumors (**Fig. 3e** and **Supplementary Fig. S1l**). Thus, *Stag2* inactivation modifies the overall tumor suppressor landscape in lung cancer.

Based on our analysis of DepMap data and the tumor-suppressive effects of STAG2, PAXIP1, and PAGR1 *in vivo,* we hypothesized that they may cooperate to suppress lung tumorigenesis. Using Tuba-seq^Ultra^, we found that inactivating *Paxip1* or *Parg1* increased the growth of *Stag2-*proficient tumors but had no impact on the growth of *Stag2*-deficient tumors (**Fig. 3f** and **Supplementary Fig. S1m**). This epistasis suggests that PAXIP1/PAGR1 and STAG2-cohesin suppress lung tumorigenesis through partially overlapping the mechanisms.

Finally, *Stag1* was the only cohesin complex component for which inactivation had a significantly different effect in *Stag2*-proficient versus *Stag2*-deficient lung tumors (**Fig. 3c,g**). *Stag1* inactivation was significantly more deleterious for *Stag2*-deficient lung tumors with effects on tumor number approaching that of inactivating the known essential gene *Pole* (**Fig. 3g** and **Supplementary Fig. S1n)**. These results are consistent with the well-established synthetic lethality of Stag1 and Stag2 and the essential role of cohesin in general in this tumor type (4,43). Collectively, these quantitative analyses demonstrate that STAG2 inactivation impacts the overall tumor suppression landscape of lung cancer, is synthetic lethal with STAG1, and is epistatic to the PAXIP1/PAGR1 complex.

### STAG2-deficient cancer cells have gene expression changes related to cell differentiation, metabolic pathways, and response to immune cell infiltration

To characterize the molecular effects of STAG2-deficiency, we initiated tumors in *KT* and *KT;Stag2^flox^* mice and used FACS to isolate neoplastic cells (DAPI^neg^, lineage^neg^, Tomato^pos^ cells) for molecular analyses (**Fig. 4a**) (44). To capture any changes in the molecular output of STAG2 deficiency during tumor development, we collected samples after 8 and 16 weeks of tumor growth. We initiated tumors using different viral titers such that mice would have similar overall tumor burden at time of analysis (**Fig. 4a** and **Supplementary Fig. S2a-b**).

**Figure 4.**
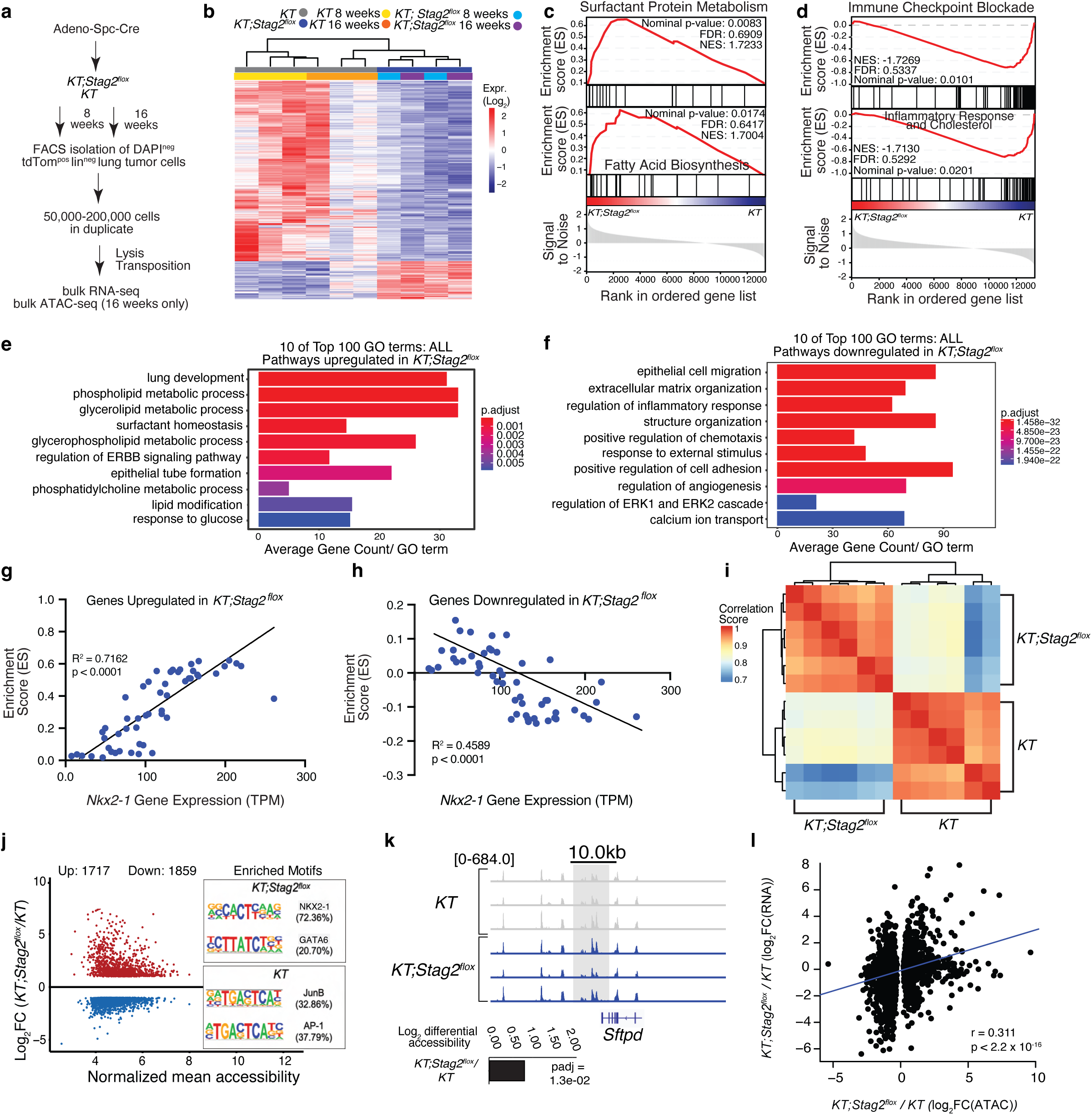
STAG2 inactivation increases tumor-related metabolic processes and cell differentiation. **a)** Schematic of tumor initiation in *KT* and *KT;Stag2^flox^* mice. Mice grew tumors until 8 weeks to represent large atypical adenomatous hyperplasia and small adenomas and 16 weeks to represent larger solid adenomas and early adenocarcinomas (Marjanovic *et al.*, 2020). Outline of tumor cell sorting and sample preparation for bulk RNA-seq and bulk ATAC-seq. **b)** Upregulated and downregulated genes (n = 2545 genes) in *KT* relative to *KT;Stag2^flox^*tumors (absolute value of log_2_ fold change (|log_2_FC|) > 1, FDR < 0.01). **c-d)** Gene Set Enrichment Analysis (GSEA) pathways enriched in *KT;Stag2^flox^* relative to *KT* tumors **(c)** and enriched in *KT* relative to *KT;Stag2^flox^* tumors **(d)**. **e-f)** GO Term Gene Count Analysis with ClusterProfiler and EMBL-EBI GO:Term Category Analysis established pathways enriched from upregulated genes for *KT;Stag2^flox^*/*KT* mice **(e)** and from down-regulated genes for *KT;Stag2^flox^*/*KT* mice **(f)**. **g-h)** ES versus NKX2-1 gene expression (TPM) from Single Sample Gene Set Enrichment Analysis (ssGSEA) for *KPT* and *KT* samples (Chuang *et al.*, 2017) for gene set from genes upregulated in *KT;Stag2^flox^* versus *KT* **(g)** and genes downregulated in *KT;Stag2^flox^* versus *KT* **(h)**. **i)** Rank correlation of chromatin accessibility across *KT* and *KT;Stag2^flox^* samples. Samples cluster into two distinct groups. **j)** Differential accessibility across 3576 significant peaks in *KT* and *KT;Stag2^flox^*mice (3 mice/group). The x-axis represents the log_2_ mean accessibility per peak and the y-axis represents the log_2_ fold change in accessibility. Colored points are significant (|log_2_FC| > 1,FDR < 0.05). Red points are increased chromatin accessibility in *KT;Stag2^flox^* accompanied by transcription factor hypergeometric motif enrichment in *KT;Stag2^flox^*, and blue points are decreased chromatin accessibility in *KT;Stag2^flox^* accompanied by transcription factor hypergeometric motif enrichment in *KT* tumors. **k)** ATAC-seq signal tracks for the *Sftpd* gene locus in tumor from *KT* and *KT;Stag2^flox^* mice. **l)** Comparison of log_2_ fold change for *KT* and *KT;Stag2^flox^* mice for top genes from both bulk RNA-seq and bulk ATAC-seq. r = 0.311, p < 2.2 × 10-16.

*Stag2* inactivation in these samples was validated by mapping reads to the floxed exon of *Stag2* and measuring *Stag2* expression (**Supplementary Fig. S2c-d**). Based on hierarchical clustering and principle component analysis (PCA), samples from *KT* and *KT;Stag2^flox^*mice formed distinct clusters (**Fig. 4b** and **Supplementary Fig. S2e**). Comparisons between neoplastic cells from all *KT* and *KT;Stag2^flox^* samples uncovered >2500 differentially expressed genes (absolute value of log_2_ fold change (|log_2_FC|) > 1, FDR <0.01; **Fig. 4b** and **Supplementary Table S5**). Using Gene set enrichment analysis (GSEA) and Gene Ontology (GO) term enrichment, we uncovered upregulation of pathways related to surfactant protein metabolism (exemplified by significantly increased expression of the transcription factor *Nkx2-1* and canonical NKX2-1-regulated surfactant protein genes), lung development, and several metabolic processes in *Stag2*-deficient neoplastic cells (**Fig. 4c,e** and **Supplementary Fig. S2f-g**). Consistent with this gene expression data, immunohistochemical staining for NKX2-1 showed that *Stag2*-deficient tumors were significantly more likely to have high expression for NKX2-1 than *Stag2*-proficient tumors (p < 0.001; **Supplementary Fig. S2h**). Significantly down regulated gene sets in *Stag2*-deficient neoplastic cells were related to epithelial response to immune cells and extracellular matrix organization (**Fig. 4d,f**). Across an independent dataset of lung tumorigenesis in oncogenic KRAS-driven models, *Nkx2-1* expression positively correlated with expression of genes that were upregulated by STAG2-deficiency (R^2^ = 0.7162, p < 0.0001) and negatively correlated with expression of genes that were downregulated by STAG2 inactivation (R^2^ = 0.4589, p < 0.0001) consistent with NKX2-1 driving some of these differences (**Fig. 4g-h**) (45).

To assess whether Stag2 deficiency also impacts the underlying chromatin state, we performed ATAC-seq on neoplastic cells isolated from *KT* and *KT;Stag2^flox^* mice. Neoplastic cells from *KT* and *KT;Stag2^flox^* mice clustered separately and had comparable library quality (**Fig. 4i, Supplementary Fig. S2i** and **Methods**). Stag2-deficient neoplastic cells have several thousand regions of increased and decreased chromatin accessibility relative to Stag2-proficient neoplastic cells (|log_2_FC| > 1, FDR < 0.05) (**Fig. 4j, Supplementary Fig. S2j,** and **Supplementary Table S9**). Interestingly, regions with increased accessibility in *Stag2*-deficient neoplastic cells were enriched for NKX2-1 and GATA family transcription factor motifs, which are well-established regulators of lung-specific lineage programs and lung development (**Fig. 4j** and **Supplementary Fig. S2k**) (45–48). Additionally, there were regions with increased accessibility proximal to lung lineage genes, such as *Sftpd* (**Fig. 4k**). Regions with decreased accessibility in *Stag2-*deficient neoplastic cells were enriched for JUNB, AP-1, FOS, BATF, and FRA1/2 motifs (**Fig. 4j** and **Supplementary Fig. S2l**). Notably, the gene expression and chromatin accessibility changes in neoplastic cells from *KT* and *KT;Stag2^flox^* mice were positively correlated (R = 0.311, p < 2.2 × 10^-16^) (**Fig. 4l**). Collectively, these data show that STAG2 inactivation rewires the epigenome and changes gene expression states.

### STAG2 inactivation results in differential DNA looping that correlates with increased gene expression

Cohesin regulates 3D genome structure which can facilitate enhancer-promoter interactions and impact gene expression (2,49). Thus, to determine whether STAG2 inactivation impacts chromatin looping, we performed chromosome conformation capture (Hi-C) on neoplastic cells isolated from *KT* and *KT;Stag2^flox^* mice (**Fig. 5a** and **Supplementary Fig. 3a**). *Stag2*-deficient neoplastic cells gained and lost DNA loops compared to *Stag2*-proficient cells. Specifically, in addition to > 7,500 common loops there were ∼3,000 loops that were lost/reduced in Stag2-deficient cells (*KT* unique loops) and >5,700 loops that were gained/stronger in *Stag2-* deficient cells (*KT;Stag2^flox^* unique loops; **Fig. 5b-c**). Furthermore, *KT;Stag2^flox^* unique loops were on average larger (p = 5.5 × 10^-28^) than *KT* unique loops (**Fig. 5d** and **Supplementary Fig. S3b**). We identified unique and shared loop “anchors” between *KT* and *KT;Stag2^flox^* samples. Notably, many of the unique loops share one anchor site with loops in the corresponding *KT* or *KT;Stag2^flox^* samples **(Fig. 5e)**. On average the loop anchors were more distal in *KT;Stag2^flox^* samples compared with *KT* suggesting loop extrusion may favor different anchor sites in the absence of STAG2 **(Supplementary Fig. S3c**).

**Figure 5.**
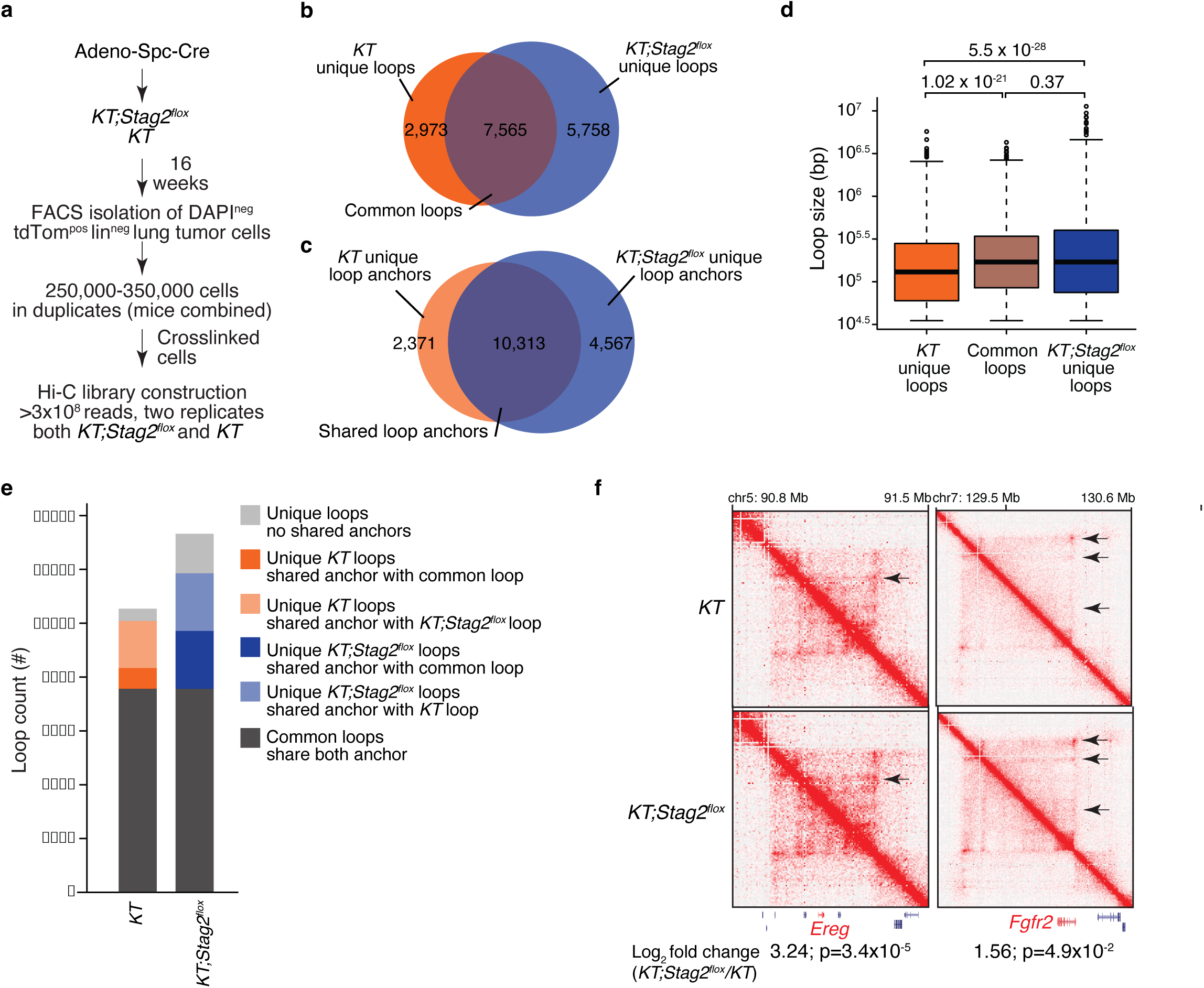
*Stag2* deficiency impacts overall chromatin looping with effects on gene expression. **a)** Schematic of tumor initiation with Adeno-Spc-Cre in *KT* and *KT;Stag2^flox^* mice. Outline of tumor cell sorting, crosslinking, and sample preparation for Hi-C. **b)** Venn diagram of loops in *KT* and *KT;Stag2^flox^* samples. **c)** Venn diagram of loop anchors in *KT* and *KT;Stag2^flox^* samples. **d)** Sizes for *KT* unique loops, common loops, and *KT;Stag2^flox^* unique loops. Boxes show median +/− interquartile range. Whiskers show standard error. P-values (Wilcoxon rank test) are shown. **e)** Loops in *KT* and *KT;Stag2^flox^* samples were compared and sorted into indicated categories based on their unique and shared loop anchors. **f)** Differences in chromatin looping and gene expression (from RNA-seq data, log_2_ fold change (*KT;Stag2^flox^*/*KT*); p-value) for *Ereg* and *Fgfr2*.

To better understand whether the differences in 3D chromatin looping contribute to the molecular changes caused by *Stag2* inactivation, we assessed whether the anchor sites of unique loops are associated with transcription start sites (TSSs) of differentially expressed genes. By overlapping differentially expressed genes with the coordinates of the anchor sites of unique loops, we found that genes upregulated in *KT;Stag2^flox^* samples are enriched proximal to the anchor sites of *KT;Stag2^flox^* unique loops (*i.e.* the gene TSS is within the anchor site) (85 genes, p = 5.3 × 10^-8^) (**Supplementary Fig. S3d**). Similarly, genes with higher expression in *KT* samples were enriched proximal to anchor sites of *KT* unique loops (67 genes, p = 5.6 × 10^-7^) (**Supplementary Fig. S3e** and **Supplementary Table 11**). For differentially expressed genes that are associated with unique anchors sites in either *KT* or *KT;Stag2^flox^* samples, we analyzed whether the corresponding anchor (*i.e.* the other side of the unique loop) had changes in chromatin accessibility, which might occur if the generation of novel enhancers led to the formation of the differential chromatin interactions. Interestingly, >75% of the corresponding anchor sites did not significantly change in chromatin accessibility. This suggests that the changes in 3D chromatin looping are the result of STAG2 inactivation not because of enhancer re-programming. Additionally, many of the genes with increased expression and proximity to unique anchor sites in *Stag2*-deficient neoplastic cells, such as EREG and FGFR, are associated with tumor growth pathways (**Fig. 5f**). We also found two loci associated with unique loops in *Stag2*-deficient cells, that have multiple genes that are all upregulated, possible due to the different loop extrusion in the absence of STAG2 (**Supplementary Fig. S3f-g**). These regions include genes implicated in lung lineage development and lung adenocarcinoma and progression (50–52). Altogether, this genome conformation data strongly suggests that STAG2 inactivation leads to changes in 3D chromatin looping and activation of downstream pathways that could promote tumor growth and alter differentiation.

### PAXIP1 and STAG2-cohesin control conserved gene expression programs

As our genetic epistasis data suggest that the tumor-suppressive function of PAXIP1 and STAG2-cohesin are related, we next determined the extent to which the molecular programs driven by PAXIP1 and STAG2-cohesin overlap. To characterize the molecular effects of PAXIP1 deficiency in lung tumors, neoplastic cells were FACS-isolated from tumors initiated in *KT;H11^LSL-Cas9^* mice with Lenti-sg*Inert*/Cre or Lenti-sg*Paxip1*/Cre (**Fig. 6a**). We performed bulk RNA-seq on 3-4 samples from each group (hereafter sg*Inert* and sg*Paxip1*). Samples cluster based on their genotype (**Fig. 6b** and **Supplementary Fig. S4a**). Many (∼540) genes were differentially expressed between sg*Paxip1* and sg*Inert* neoplastic cells (|log_2_FC| > 1 and FDR < 0.01) (**Fig. 6b** and **Supplementary Table S6**).

**Figure 6.**
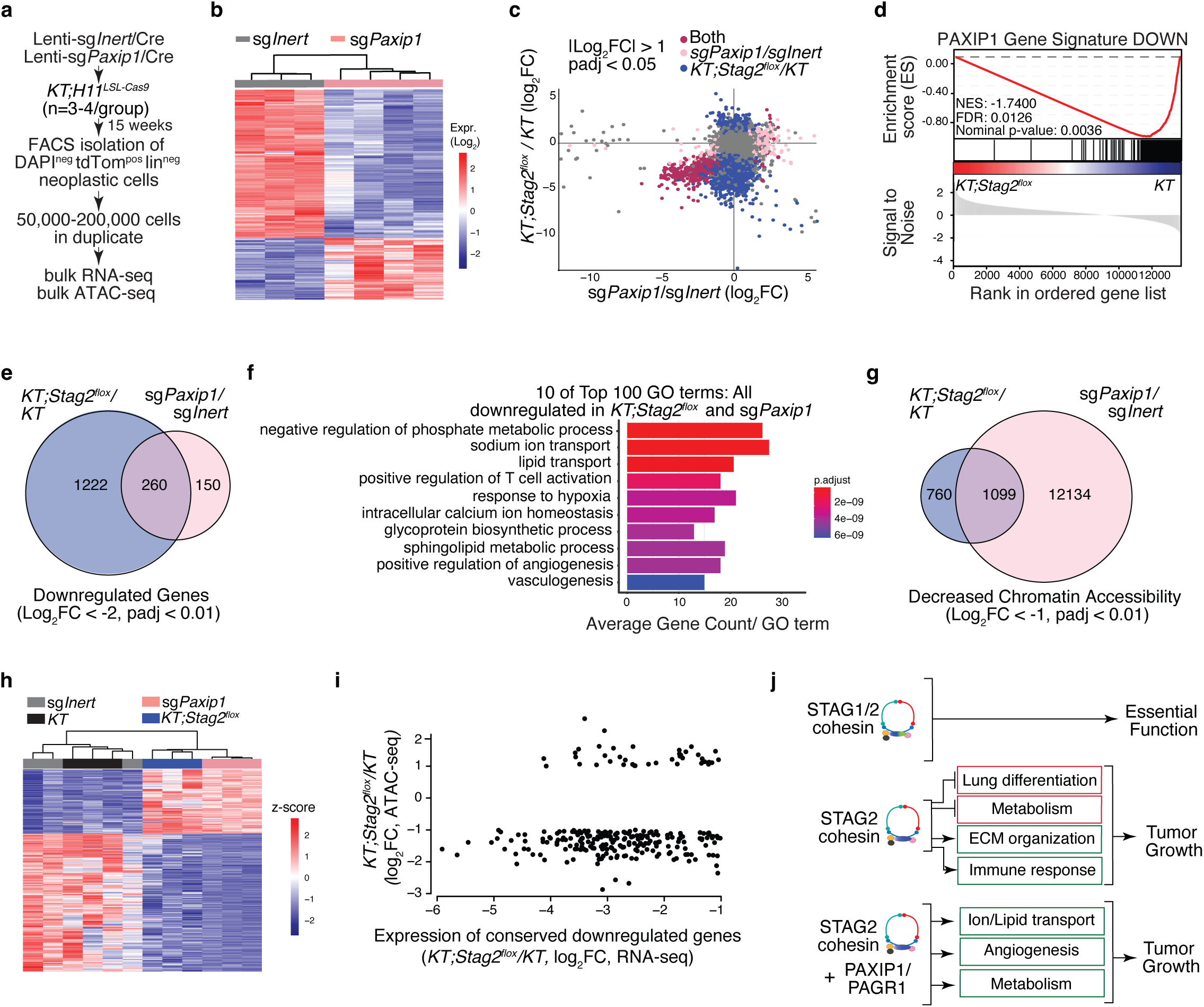
PAXIP1 and STAG2-cohesin mechanisms of tumor suppression are conserved. **a)** Schematic of tumor initiation with Lenti-sg*Paxip1*/Cre or Lenti-sg*Inert*/Cre in *KT;H11^LSL-Cas9^* mice (3-4 mice/group). Mice grew tumors for 15 weeks. Outline of tumor cell sorting and sample preparation for bulk RNA-seq and bulk ATAC-seq. **b)** Upregulated and downregulated genes (n=540 genes) in *KT;H11^LSL-Cas9^* sg*Inert* relative to *KT;H11^LSL-Cas9^* sg*Paxip1* tumors (|log_2_FC| > 1, padj <0.01). **c)** Positive correlation between *KT;Stag2^flox^/KT* (log_2_FC) and *KT;H11^LSL-Cas9^* sg*Paxip1*/*KT;H11^LSL-Cas9^* sg*Inert* (log_2_FC). Each dot is a gene. Dots |log_2_FC| > 1, padj <0.05 in both are maroon, dots |log_2_FC| > 1, padj <0.05 in *KT;H11^LSL-Cas9^* sg*Paxip1*/*KT;H11^LSL-Cas9^* only are pink, and dots |log_2_FC| > 1, padj <0.05 in *KT;Stag2^flox^/KT* are dark blue. Dots |log_2_FC| > 1, padj <0.05 in neither are grey. **d)** Downregulated PAXIP1 gene signature enrichment in rank-ordered gene list for *KT* versus *KT;Stag2^flox^*. **e)** Venn diagram of shared downregulated genes (log_2_FC < −2, padj <0.01) between *KT;Stag2^flox^/KT* and *KT;H11^LSL-Cas9^* sg*Paxip1*/*KT;H11^LSL-Cas9^* sg*Inert*. **f)** GO Term Gene Count Analysis with ClusterProfiler and EMBL-EBI GO:Term Category Analysis established from conserved downregulated pathways between *KT;Stag2^flox^/KT* and *KT;H11^LSL-Cas9^* sg*Paxip1*/*KT;H11^LSL-Cas9^* sg*Inert*. **g)** Venn diagram of shared regions of decreased chromatin accessibility (log_2_FC < −1, padj < 0.01) between *KT;Stag2^flox^/KT* and *KT;H11^LSL-Cas9^* sg*Paxip1*/*KT;H11^LSL-Cas9^*sg*Inert*. **h)** Accessibility z-score for regions with significantly different accessibility in both *KT;Stag2^flox^/KT* and *KT;H11^LSL-Cas9^* sg*Paxip1*/*KT;H11^LSL-Cas9^*sg*Inert* ATAC-seq samples (|log_2_ FC| > 1, padj < 0.01) that are also associated with genes with significantly different expression in both *KT;Stag2^flox^/KT* and *KT;H11^LSL-Cas9^* sg*Paxip1*/*KT;H11^LSL-Cas9^*sg*Inert* neoplastic cells (|log_2_ FC| > 1, padj < 0.01). Each row is a differentially accessible region. **i)** Enrichment for regions with significant decreased accessibility in *KT;Stag2^flox^/KT* (|log_2_FC| > 1, padj < 0.01) samples associated with genes with significantly decreased expression in both *KT;Stag2^flox^/KT* and *KT;H11^LSL-Cas9^* sg*Paxip1*/*KT;H11^LSL-Cas9^*sg*Inert* neoplastic cells (log_2_FC < - 1, padj < 0.01). Changes in accessibility and expression in *KT;Stag2^flox^/KT* is shown. Each dot is a differentially accessible region. **i)** Model of lung tumor suppression regulated by STAG2-cohesin-PAXIP1/PAGR1 axis.

Interestingly, genes that were regulated by both STAG2 and PAXIP1 were almost exclusively downregulated in *Stag2*-deficient and *Paxip1*-deficient neoplastic cells (**Fig. 6c**). Genes that were downregulated in sg*Paxip1* neoplastic cells (“PAXIP1 DOWN”), were downregulated in *KT;Stag2^flox^* neoplastic cells (p-value < 0.0036; **Fig. 6d**). However, genes that were downregulated by *Stag2* deficiency were either equally dependent on PAXIP1 or not dependent on PAXIP1. Pathways enriched in the genes downregulated in both *KT;Stag2^flox^* and *KT;H11^LSL-Cas9^* sg*Paxip1* tumors include those related to ion transport, metabolic processes, and angiogenesis (**Fig. 6e-f**). Genes that were upregulated in *KT;H11^LSL-Cas9^* sg*Paxip1* neoplastic cells were not upregulated in *KT;Stag2^flox^* neoplastic cells and vice versa (**Supplementary Fig. S4b-g**). Given the broad changes in chromatin accessibility induced by *Stag2* deficiency, we performed ATAC-seq on sg*Inert* and sg*Paxip1* neoplastic cells (**Fig. 6a**). These samples were of similar quality and clustered by genotype (**Supplementary Fig. S4h**). There were >4200 regions with decreased chromatin accessibility and >13000 regions with increased chromatin accessibility in neoplastic cells from sg*Inert* versus sg*Paxip1* mice (|log_2_FC| > 1, FDR < 0.05) (**Supplementary Fig. S4i-j** and **Supplementary Table S9**). Motifs for NKX2-1, ZEB1, and GATA were enriched in regions with increased accessibility in sg*Paxip1* neoplastic cells, and motifs for TCF21 and BHLHA15 were enriched in regions with decreased accessibility (**Supplementary Fig. S4k-l**). Motifs for DLX1, SMAD2, and HOXA2 were enriched in regions with decreased accessibility that overlap between sg*Paxip1* neoplastic cells and STAG2-deficient cells (**Supplementary Fig. S4m**). Although there was motif enrichment for NKX2-1 in regions with increased accessibility in sg*Paxip1* neoplastic cells, neither *Nkx2-1* nor *Nkx2-1-*regulated genes were upregulated in sg*Paxip1* neoplastic cells suggesting that regulation of cell differentiation through NKX2-1 is a STAG2-specific mechanism (**Supplementary Fig. S4f-g** and **Supplementary Fig. S2f-g)**.

Similar to gene expression, there were more shared regions with decreased rather than increased chromatin accessibility (**Fig. 6g, Supplementary Fig. S4j** and **Supplementary Table S9**). Interestingly, many downregulated genes conserved between *KT;Stag2^flox^* and sg*Paxip1* had decreased accessibility in Stag2-deficient and Paxip1-deficient neoplastic cells (**Fig. 6h-i** and **Supplementary Table S7**). Lastly, there was no overlap between genes regulated by PAXIP and genes regulated by STAG2-dependent chromatin looping (**Supplementary Fig. 4n**). Collectively, these results emphasize that there are distinct PAXIP1-dependent and PAXIP1-independent STAG2-mediated tumor suppressive programs, consistent with the overall tumor-suppressive effects of PAXIP1, PAGR1 and STAG2 *in vivo* (**Fig. 6j** and **Supplementary Fig. S5**).

## DISCUSSION

Here, we coupled multiplexed and quantitative functional genomics within autochthonous lung cancer models, *in vivo* genetic epistasis experiments, and molecular analyses to uncover mechanisms of STAG2-mediated tumor suppression and link it to PAXIP1/PARG1. Tumor suppression in lung cancer was specific to STAG2-cohesin rather than STAG1-cohesin or overall cohesin. Cohesin components beyond *STAG2* are also frequently mutated in human lung cancer, and our results show that heterozygous inactivation of a core cohesin gene increased tumor growth. We show that SMC3 exhibits both essential and haploinsufficient tumor suppressor gene characteristics, further emphasizing the versatility and power of *STAG2*-cohesin as a tumor suppressor. Thus, heterozygous or hypomorphic mutations in core or auxiliary cohesin components likely reduce STAG2-cohesin-mediated tumor suppression and make the fraction of human lung tumors that are driven by this mechanism greater than only those with *STAG2* mutations (**Fig. 1** and **Supplementary Fig. S2**).

Using cancer cell line dependency data, multiple studies have revealed a correlation between the dependency scores for *STAG2* and the *PAXIP1-PAGR1* complex (8,42), which we show extends to cell lines driven by oncogenic KRAS as well as to the subset of those cell lines derived from lung adenocarcinoma (**Fig. 2b-c**). Previous studies using cancer cell lines have shown that inactivation of STAG2 and PAXIP1 reduces growth, leading to the suggestion that these genes promote cancer cell growth (42,53). However, our functional data show that STAG2, PAXIP1, and PAGR1 normally constrain lung tumor growth *in vivo*, consistent with many studies that document STAG2 as a tumor suppressor (9,11–14,19,30). Thus, while cell lines may correctly uncover correlated gene effects, their optimized growth in culture and/or lack of physiologic context limit their ability to identify functional tumor suppressors.

To identify genes that cooperate with STAG2 to suppress tumor growth *in vivo*, we compared the tumor-suppressive effects of a broad panel of genes in STAG2-proficient and - deficient autochthonous lung tumors (**Fig. 3**). These data uncovered relatively few genes whose effects were impacted by *Stag2*-deficiency, emphasizing the uniqueness of the interaction between STAG2 and PAXIP1/PAGR1 (**Fig. 3c)**. In this experiment, we did not address the role of inflammation in KRAS-driven tumor progression, which could serve as a critical dimension for future studies (54,55). PAXIP1 and PAGR1 have been implicated in several biological processes, including DNA damage responses (26,27,29,56). Thus, we initially considered whether the reduced impact of *Paxip1* and *Pagr1* inactivation on tumor growth relative to *Stag2* inactivation could be due to partially offsetting tumor-suppressive and essential DNA damage functions (**Fig. 3f** and **Supplementary Fig. S5a-b**). However, *Paxip1* or *Pagr1* inactivation in the context of *Stag2*-deficiency (*i.e.,* in *KT;H11^LSL-Cas9^;Stag2^flox^* mice) did not reduce tumorigenesis, as would be expected if these two genes had a second essential function (**Fig. 3f** and **Supplementary Fig. S5a-b**). Furthermore, PAXIP1/PAGR1 are required for the expression of a subset of STAG2-regulated genes, some of which are likely responsible for PAXIP1/PAGR1-mediated tumor suppression **(Fig. 5** and **Supplementary Fig. S5a-b**).

STAG2-mediated tumor suppression is also mediated by PAXIP1/PAGR1-independent gene expression programs (**Fig. 5** and **Supplementary Fig. S3**). Interestingly, in contrast to many genetic alterations that increase tumorigenesis while decreasing or changing differentiation, STAG2-deficient tumors express higher levels of the lung lineage defining transcription factor NKX2-1 as well as canonical genes related to ATII differentiation (**Fig. 4g-h, j-k** and **Supplementary Fig. S2-3**). Additionally, there is increased DNA loop formation in STAG2-deficient tumors at loci that overlap with genes involved in lung differentiation and tumorigenesis, such as genes related to ATII differentiation and the EGF pathway (**Fig. 6d** and **Supplementary Fig. S4e-g**). Thus, it is interesting to note the potential stabilization of this cellular state by STAG2 deficiency, which otherwise might be anticipated to be selected against (57). Genomic regions with increased accessibility in STAG2-deficient tumors are also enriched for NKX2-1 binding sites (**Fig. 4k**). Whether NKX2-1 interacts directly or indirectly with STAG2-cohesin-PAXIP1/PAGR1 and whether an NKX2-1-regulated oncogenic programs drive growth in this context will be important areas of future investigation (58).

While we show clear evidence for a genetic interaction between STAG2-cohesin and the PAXIP1/PAGR1 complex during lung tumor suppression this data does not distinguish whether PAXIP/PAGR1 functions upstream or downstream to STAG2-cohesin (**Fig. 3f**). Our data and existing literature on PAXIP1/PAGR1 and STAG2-cohesin are most consistent with a model in which PAXIP1/PAGR1 is initially recruited to genome loci (perhaps through direct interactions with lineage-specific transcription factors), followed by STAG2-cohesin localization at those regions, and the formation of genomic contacts that ultimately impact gene expression to suppress lung tumor growth (42,59). A direct interaction between PAXIP1/PAGR1 and STAG2-cohesin is only weakly supported by existing mass spectrometry data on STAG2-, cohesin-, and PAXIP1-interacting proteins (25,60). PAXIP/PAGR1 and STAG2-cohesin could interact weakly or transiently, through an intermediate protein, or be functionally linked through a less direct mechanism such as PAXIP1/PAGR1-mediated recruitment of histone modifying enzymes followed by preferential localization of STAG2-cohesin at these regions (28,61). Interestingly, while our Hi-C data uncovered unique loops that overlap with genes impacted by STAG2 inactivation, these unique loops did not overlap with genes impacted by both STAG2 inactivation and PAXIP1 inactivation (**Supplementary Fig. S4n** and **Supplementary Table 7, 11**). Further biochemical, genetic, and genomic experiments including CTCF ChIA-PET and CUT&RUN should further clarify the interaction between STAG2-cohesin and the PAXIP1/PAGR1 complex as well as their genomic localization during tumor suppression.

Interestingly, the tumor-suppressive STAG2-PAXIP1/PAGR1 axis may transcend cancer type. In human cell lines, *STAG2, PAXIP1,* and *PAGR1* have highly correlated effects in bladder cancer, Ewing’s sarcoma, acute myeloid leukemia (AML), and lung adenocarcinoma, all of which also have frequent mutations in *STAG2* (**Supplementary Fig. S1f-i**) (31–33). In particular, the correlation of *STAG2* with *PAXIP1* and *PAGR1* in AML was strikingly high (r = 0.86 and r=0.90, respectively) (**Supplementary Fig. S1f-g**). In AML, *STAG2* mutations and heterozygous mutations in *SMC3* are well-established regulators of the pre-leukemic state (6). Interestingly, cancer sequencing studies have also identified mutations in *PAXIP1* and *PAGR1* in AML and down regulation of *PAXIP1* expression in AML (**Supplementary Fig. S5c-e**) (31–33,62). Thus, the PAXIP1/PAGR1 complex may represent an important unexplored driver in AML.

Collectively, our data not only highlight a STAG2-PAXIP1/PAGR1 tumor-suppressive axis that may transcend cancer types but also establishes distinct PAXIP1-independent and PAXIP1-dependent STAG2-mediated tumor-suppressive functions. Our findings underscore the important roles of STAG2-cohesin in suppressing lung tumorigenesis.

## ACKNOWLEDGEMENTS

We thank the Stanford Shared FACS Facility for flow cytometry and cell sorting services, the Stanford Veterinary Animal Care Staff for expert animal care, Human Pathology/Histology Service Center, and Stanford Protein and Nucleic Acids Facility. We would like to acknowledge the American Association for Cancer Research and its financial and material support in the development of the AACR Project GENIE registry, as well as members of the consortium for their commitment to data sharing. Interpretations are the responsibility of study authors. We thank D. Solomon, D. Hargreaves, J. Sage, L. Attardi, P. Khavari, J. Carette, P. Du, B. Chick, N. Avina Ochoa, and members of the Winslow and Crabtree laboratories for helpful comments and experimental support. We thank A. Melnick for providing *Smc3^floxed^*mice. E.L.A. was supported by the HHMI Gilliam Fellowship for Advanced Study (GT14928) and a National Cancer Institute Predoctoral to Postdoctoral Fellow Transition Award (F99CA284289). Y.J.T. was supported by the Canadian Institute of Health Research (CIHR) postdoctoral fellowship (MFE-176568). K.L.H. was supported by a Stanford Graduate Fellowship and a National Cancer Institute Predoctoral to Postdoctoral Fellow Transition Award (F99CA274692). J.B. was supported by the HHMI Hanna H. Gray Fellowship Award. H.C. was supported by a Tobacco-Related Disease Research Program (TRDRP) fellowship (28FT-0019). D.N.D. was supported by the National Cancer Institute Predoctoral to Postdoctoral Fellow Transition Award (K00-CA245784). J.D.H was supported by an American Cancer Society postdoctoral fellowship (PF-21-112-01-MM) and a Tobacco-Related Disease Research Program (TRDRP) fellowship (T31FT1619). P.A.R was supported by the National Science Foundation Graduate Research Fellowship Program (DGE-2146755) and the Stanford Graduate Fellowship. This work was supported by NIH R01-CA231253 (to M.M.W) and R01-CA234349 (to M.M.W and D.A.P) and in part by the Stanford Cancer institute support grant (NIH P30-CA124435).

## CONTRIBUTIONS

E.L.A. and M.M.W. conceptualized and designed the study. Y.J.T designed the Tuba-seq^Ultra^ methodology. E.L.A., Y.J.T., and L.A. prepared Tuba-seq libraries. E.L.A., H.X., J.B., Z.X., and K.L.H. designed the computational pipelines, methodology, and performed formal analysis. E.L.A., M.M.W., Z.X., and H.X. wrote the manuscript with comments from all authors. E.L.A., J.D.H., S.L., D.N.D., Z.X., J.R.D., M.M.W. and K.L.H. edited the manuscript. E.L.A., Y.J.T., H.C., S.L., D.N.D., P.R., R.L., and L.A. acquired data. M.M.W., H.Y.C., J.R.D., and D.A.P. provided resources.

## DECLARATION OF INTERESTS

M.M.W and D.A.P are founders and hold equity in Guide Oncology. H.Y.C. is a co-founder of Accent Therapeutics, Boundless Bio, Cartography Biosciences, 428 Orbital Therapeutics, and an advisor of 10x Genomics, Arsenal Biosciences, Chroma 429 Medicine, and Spring Discovery.

## METHODS

### Design, generation, barcoding, and production of lentiviral and adenoviral vectors

sgRNA sequences for each putative tumor suppressor gene from the Tuba-seq^Ultra^ pools were chosen utilizing a combination of CRISPick (https://portals.broadinstitute.org/gppx/crispick/public) and the Brie library (63). Firstly, the sgRNAs from the Brie library were filtered based on (1) their predicted on-target efficacy score (score > 0.2^2^) from CRISPick and (2) off-target risk. To evaluate off-target risk, the sgRNA sequence + NGG were aligned to mouse reference genome assembly (GRCm39). sgRNAs that have a perfect match or a tolerable mismatch (a single mismatch occurring at a position located more than 10 bases away from the PAM sequence) in an off-target location were regarded as high off-target risk and were excluded. For the Brie sgRNAs that successfully passed this filter, a maximum of two sgRNAs were selected based on their on-target efficacy score. To ensure the inclusion of three sgRNAs per gene, additional sgRNAs were designed using CRISPick, adhering to the aforementioned two criteria. This approach resulted in a final set of sgRNAs, comprising both tested sgRNAs from the Brie library and novel sgRNAs designed based on one of the most reliable sgRNA-predicting algorithms available (64). All sgRNA sequences used are shown in **Supplementary Tables 1, 2, 4**.

To generate vectors for Tuba-seq^Ultra^, barcode-containing oligo pools were cloned at the 3’ end of the bU6 promoter in the Tuba-seq^Ultra^ backbone vector by Gibson assembly and sgRNA libraries were added using Golden Gate assembly (35). Prior to adding the barcoded oligo pools, the last 20 bp of the bU6 promoter and 137 bp filler from the Tuba-seq^Ultra^ backbone vector were removed, followed by DpnI digestion to remove remaining methylated parental vector. The single-stranded barcoded oligos were cloned into the linearized and Dpn1-digested Tuba-seq^Ultra^ backbone vector with Gibson assembly to diversify the sequence of the last 20 bp of the bU6 promoter. PCR-amplified sgRNA pools (TWIST Biosciences) were cloned into the BsmBI-v2-digested barcoded Tuba-seq^Ultra^ vector pool using Golden Gate assembly. The products were transformed into NEB 10-β electrocompetent bacteria and several million colonies were pooled, followed by plasmid extractions (Qiagen Plasmid Plus Midiprep kit) and sequencing (Novogene Corporation, Inc.) to determine the barcode and sgRNA representation (**Supplementary Fig. S2a**). To generate Lenti-sgRNA/Cre vectors encoding individual sgRNAs, individual sgRNAs were cloned downstream of the U6 promoter in a pLL3.3 backbone containing the PGK promoter and Cre using site-directed mutagenesis (34).

Lentiviral vectors were produced using polyethylenimine (PEI)-based transfection of 293T cells with delta8.2 and VSV-G packaging plasmids in 150 mm cell culture plates. Sodium butyrate (Sigma Aldrich, B5887) was added 8 hours after transfection to achieve a final concentration of 20 mM. Medium was refreshed 24 hours after transfection. 20 mL of virus-containing supernatant was collected 36 and 48 hours after transfection. The two collections were then pooled and concentrated by ultracentrifugation (25,000 rpm for 1.5 hours), resuspended overnight in 100 µL PBS, and frozen at −80°C. Adeno-Spc-Cre was purchased from University of Iowa Core Web.

We generated a pool of barcoded lentiviral vectors encoding Cre along with sgRNAs targeting a panel of genes. Vectors with sgRNAs targeting essential genes (**Supplementary Table S1,2, and 4)** and ‘inert’ sgRNAs were also generated. In this lentiviral system, a diverse barcode is integrated into the bovine U6 (bU6) promoter directly 5’ of the sgRNA. As a result, each clonal tumor is uniquely identified by a barcode-sgRNA element, and the sgRNA sequence indicates the specific gene targeted in each tumor.

### Mice and tumor initiation

The use of mice for the current study has been approved by the Institutional Animal Care and Use Committee at Stanford University, protocol number 26696. *Kras^LSL-G12D/+^* (RRID:IMSR_JAX:008179), *R26^LSL-tdTomato^*(RRID:IMSR_JAX:007909), *H11^LSL-Cas9^* (RRID:IMSR_JAX:027632), *Stag2^flox^*(RRID:IMSR_JAX:030902), and *Smc3^flox^* (RRID:IMSR_JAX:030559) mice have been previously described (19,65,66). All mice were on a C57BL/6 background. Lung tumors were initiated by intratracheal delivery of Lenti-sgRNA/Cre vectors or Adeno-Spc-Cre vectors (19,64). Adenoviral vectors were CaCl_2_-precipitated prior to delivery.

In our initial screen for cohesin complex members, which was part of a larger screen of regulators of lung tumorigenesis (35), cohorts of *KT* (n=14) and *KT;H11^LSL-Cas9^* (n=68) were transduced with 3 × 10^5^ and 1 × 10^5^ infectious units (IFU) per mouse, respectively (**Fig. 1a**). To generate lung tumors that were heterozygous mutant for cohesin, tumors were initiated in *KT* and *KT;Smc3^flox/wt^* mice (n=13-15 mice/group) with 1 × 10^9^ IFU Adeno-Spc-Cre (**Fig. 1f**). For our screen that included sg*Stag2*, sg*Paxip1*, and sg*Pagr1*, which was part of a larger screen of regulators of lung tumorigenesis, tumor cohorts of *KT* (n=5) and *KT;H11^LSL-Cas9^* (n=31) (35) were transduced with 3 × 10^5^ IFU and 1 × 10^5^ IFU, respectively, of the Lenti-U6^BC^sgRNA/Cre pool. To generate tumors with inactivation of PAXIP1 for RNA-seq and ATAC-seq, cohorts of *KT* (n=13) and *KT;H11^LSL-Cas9^*(n=32) mice were transduced with Lenti-sg*Paxip1*/Cre and Lenti-sg*Inert*/Cre at 1.5 × 10^5^ IFU and 3 × 10^5^ IFU per mouse, respectively (**Fig. 2d**). For our Tuba-seq^Ultra^ screen of STAG2-mediated tumor suppression, cohorts of *KT* (n=12), *KT;H11^LSL-Cas9^*(n=33), and *KT;H11^LSL-Cas9^;Stag2^flox^* (n=35) mice were transduced with 3 × 10^5^ IFU, 1 × 10^5^ IFU, and 5 × 10^4^ IFU of Lenti-U6^BC^-sgRNA/Cre per mouse, respectively (**Fig. 3a-b**). To generate lung tumors for RNA-seq and ATAC-seq, tumors were initiated in *KT* (n=20) with 5 × 10^9^ and 1 × 10^9^ IFU per mouse for collection at the 8- and 16-week timepoints, respectively, and in *KT;Stag2^flox^* (n=20) with 2.5 × 10^8^ and 2.5 × 10^7^ IFU per mouse for the 8- and 16-week timepoints, respectively (**Fig. 4a**).

### Tuba-seq^Ultra^ library generation

Genomic DNA was isolated from bulk tumor-bearing lung tissue from each mouse as previously described (19,35,67,68). Following homogenization and overnight proteinase K digestion, genomic DNA was extracted from the lung lysates using standard phenol-chloroform methods. Subsequently, Q5 High-Fidelity 2x Master Mix (New England Biolabs, M0494X) was used to amplify the U6-BC-sgRNA region from 32 μg of genomic DNA in a total reaction volume of 800 μL per sample. The unique dual-indexed primers used were Forward: AAT GAT ACG GCG ACC ACC GAG ATC TAC AC-8 nucleotides for i5 index-ACA CTC TTT CCC TAC ACG ACG CTC TTC CGA TCT-6 to 9 random nucleotides for increased diversity-GCG CAC GTC TGC CGC GCT G and Reverse: CAA GCA GAA GAC GGC ATA CGA GAT-6 nucleotides for i7 index-GTG ACT GGA GTT CAG ACG TGT GCT CTT CCG ATC T-9 to 6 random nucleotides for increased diversity-CAG GTT CTT GCG AAC CTC AT. The PCR products were purified with Agencourt AMPure XP beads (Beckman Coulter, A63881) using a double size selection protocol. The concentration and quality of the purified libraries were determined using the Agilent High Sensitivity DNA kit (Agilent Technologies, 5067-4626) on the Agilent 2100 Bioanalyzer (Agilent Technologies, G2939BA). The libraries were pooled based on lung weight, cleaned up using AMPure XP beads, and sequenced (read length 2×150bp) on the Illumina HiSeq 2500 or NextSeq 500 platform (Novogene Corporation, Inc.).

### Processing of paired-end reads to identify the U6-barcode and sgRNA

Paired ends were first merged using AdapterRemoval (69), and merged reads were parsed using regular expressions to identify the sgRNA sequence and clonal barcode. When identifying sgRNA sequences, we strictly required a perfect match with the designed sequences. The 14-nucleotide random barcode sequence possesses a high theoretical diversity of approximately 4^14^ (> 10^8^). In each mouse, there are typically fewer than 100 unique tumors associated with each sgRNA. This indicates that the probability of two genuine unique clonal barcodes being within a hamming distance of each other is extremely low. Consequently, when we encounter low-frequency clonal barcodes within a 1-hamming distance of high-frequency clonal barcodes, we attribute them to sequencing or PCR errors. These low-frequency barcodes were merged with barcodes of higher frequencies.

After applying these filtering steps, we converted the read counts associated with each barcode-sgRNA into absolute neoplastic cell numbers. This conversion was accomplished by normalizing the read counts to the number of reads in the “spike-in” cell lines added to each sample prior to lung lysis and DNA extraction. The median sequencing depth across all experiments was approximately 1 read per 50 cells. To perform statistical comparisons of tumor genotypes, we imposed a minimum tumor size cutoff of 300 cells.

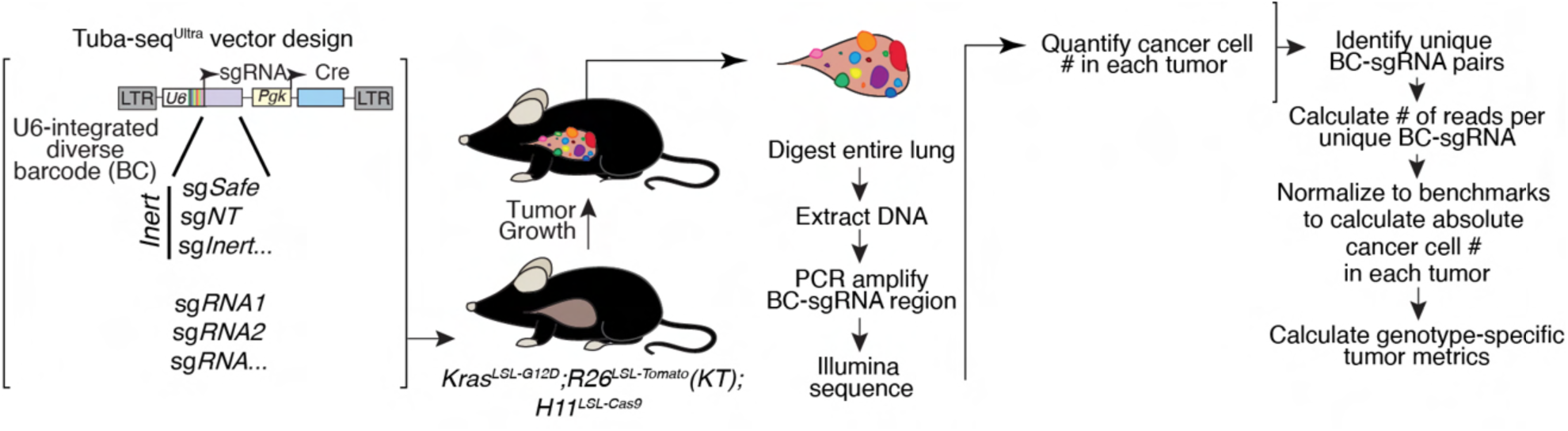
Schematic of Tuba-seq^Ultra^ vector design and tumor initiation. U6-integrated diverse barcode (U6^BC^) contains a 21-nucleotide region at the 3’ end of the bovine U6 promoter immediately downstream of the TATA box. The sgRNA-Pool/Cre consists of barcoded lenti-sgRNA/Cre vectors that contained three sgRNAs targeting each gene of interest, Safe cutting “inert” sgRNAs, and sgRNAs targeting an essential gene. *KT;H11^LSL-Cas9^* are transduced with the lenti-U6^BC^sgRNA-Pool/Cre. Fourteen-fifteen weeks after tumor initiation, we extract DNA from bulk tumor-bearing lungs and used Tuba-seq^Ultra^ to quantify the impact of targeting each regulator on tumor growth, and tumor initiation for each tumor of each genotype.

### Summary statistics for overall growth rate

To quantify the impact of each gene on tumor growth, we employed a normalization process that involved calculating statistics for tumors produced by a given sgRNA *X* (sg*X* tumors) and comparing them to the corresponding statistic of tumors generated by control sgRNAs (sg*Inert* tumors). Two key statistical measures were employed to characterize these distributions: the size of tumors at defined percentiles of the distribution (specifically the 50^th^, 70^th^, 80^th^, 90^th^, and 95^th^ percentile tumor sizes), and the log-normal mean (LN mean) size. The percentile sizes are nonparametric summary statistics of the tumor size distribution. By focusing on percentiles corresponding to the upper tail of the distribution, we specifically examined the growth of larger tumors, thereby mitigating potential confounding factors arising from variations in cutting efficiency among guides. The LN mean is the maximum-likelihood estimate of mean tumor size assuming a log-normal distribution. We normalized these two statistics calculated on tumors of each genotype to the corresponding sg*Inert* statistic. The resulting ratios reflect the growth advantage or disadvantage associated with each tumor genotype relative to the growth of sg*Inert* tumors.

For example, the relative i^th^ percentile size for tumors of genotype X was calculated as:

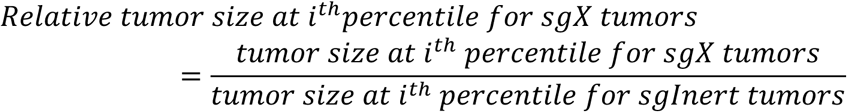

Likewise, the relative LN mean size for tumors of genotype X was calculated as:

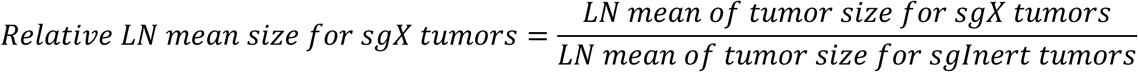

### Summary statistics for relative tumor number

In addition to the tumor size metrics described above, we characterized the effects of gene inactivation on tumorigenesis in terms of the number of tumors (“tumor number”) associated with each genotype. Unlike the aforementioned metrics of tumor size, tumor number and burden are linearly affected by lentiviral titer and are thus sensitive to underlying differences in the representation of each Lenti-sgRNA/Cre vector in the viral pool. Critically, each Tuba-seq^Ultra^ experiment included a cohort of *KT* control mice. *KT* mice lack expression of Cas9, rendering all Lenti-sgRNA/Cre vectors functionally equivalent to non-targeting sgRNA vectors in these mice. Therefore, the observed tumor number and burden associated with each sgRNA reflect their respective representation within the viral pool. This control allows us to accurately assess the specific effects of gene inactivation on tumor formation and growth, accounting for any biases introduced by the viral vector composition.

To assess the extent to which a given gene (*X)* affects tumor number, we therefore first normalized the number of sg*X* tumors in *KT;H11^LSL-Cas9^* mice by the number of sg*X* tumors in the *KT* mice:

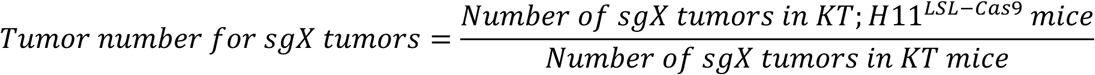

As with the tumor size metrics, we then calculated a relative tumor number by normalizing this statistic to the corresponding statistic calculated using sg*Inert* tumors:

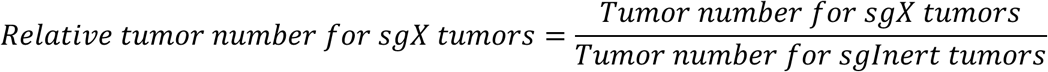

Genes that influence relative tumor number modify the probability of tumor initiation and/or the very early stages of oncogene-driven epithelial expansion, which prior work suggests are imperfectly correlated with tumor growth at later stages (44). Relative tumor number thus captures an additional and potentially important aspect of tumor suppressor gene function.

### Calculation of confidence intervals and P-values for tumor growth and number metrics

Confidence intervals and *P*-values were calculated using bootstrap resampling for each sample statistic. To account for both mouse-to-mouse and within mouse variability, we adopted a two-step, nested bootstrap approach where we first resampled mice, and then resampled tumors within each mouse to generate resampled data. 10,000 times of bootstrapping was performed to calculate 10,000 values of each statistic. 95% confidence intervals were calculated using the 2.5^th^ and 97.5^th^ percentiles of the bootstrapped statistics. Because we calculate metrics of tumor growth that are normalized to the same metrics in sg*Inert* tumors, under the null model where genotype does not affect tumor growth, the test statistic is equal to 1. Two-sided *P*-values were thus calculated as followed:

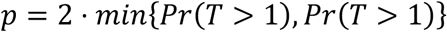

Where T is the test statistic and Pr(*T*>1) and Pr(*T*<1) were calculated empirically as the proportion of bootstrapped statistics that were more extreme than the baseline of 1. To account for multiple hypothesis testing, *P*-values were FDR-adjusted using the Benjamini-Hochberg procedure (70). Summarized statistics of all Tuba-seq^Ultra^ experiments in this study can be found in Supplementary Table 1-4.

### Analysis of DepMap data

Cancer cell line dependency data (DepMap Public 22Q2) and mutation data (Cancer Cell Line Encyclopedia) were acquired from the Broad Institute DepMap Portal (https://depmap.org/portal/). First, cell lines without non-silent mutations in *STAG2* were selected. Then the remaining cell lines were further categorized into five overlapping sets: (1) cell lines with *KRAS* mutations (with G12 or G13 substitutions), (2) cell lines derived from adenocarcinoma or non-small cell lung cancers (NSCLC), (3) cell lines obtained from bladder cancer, (4) cell lines derived from Ewing’s sarcoma, (5) cell lines derived from acute myeloid leukemia. Pearson’s and Spearman’s correlation coefficients were calculated to measure the correlation between *STAG2* and other genes using gene effect scores across the aforementioned cell line sets.

### Tumor dissociation and cancer cell sorting

Tumors were dissociated using collagenase IV, dispase, and trypsin at 37 degrees C for 30 minutes as previously described (44). Cells were stained with DAPI and antibodies against CD45 (30-F11), CD31 (390), F4/80 (BM8), and Ter119 (all from BioLegend) to exclude hematopoietic and endothelial cells. FACSAria sorters (BD Biosciences) were used for cell sorting.

### Western blotting on sorted cancer cells

Cells were lysed in RIPA buffer (50 mM Tris–HCl (pH 7.4), 150 mM NaCl, 1% Nonidet P-40 and 0.1% SDS) and incubated at 4 °C with continuous rotation for 30 min, followed by centrifugation at 12,000 rcf for 10 min. The supernatant was collected, and the protein concentration was determined by BCA assay (Thermo Fisher Scientific, 23250). Protein extracts (10–50 μg) were separated on 4–12% SDS–PAGE and transferred onto polyvinylidene fluoride membranes. The membranes were blocked with 5% non-fat milk in TBS with 0.1% Tween 20 (TBST) at room temperature for 1 h, cut according to the molecular weight of the target protein (with at least two flanking protein markers), followed by incubation with primary antibodies diluted in TBST (1:1,000) at 4 °C overnight. After three 10 min washes with TBST, the membranes were incubated with the appropriate secondary antibody conjugated to HRP diluted in TBST (1:10,000) at room temperature for 1 h. After three 10 min washes with TBST, protein expression was quantified with enhanced chemiluminescence reagents (Fisher Scientific, PI80196).

Antibodies used in this study: HSP90 (BD Biosciences, 610418), GAPDH (Cell Signaling, 5174S), PAXIP1 (Sigma-Aldrich, ABE1877), STAG2 (Santa Cruz Biotech., SC-81852), goat-anti-rabbit IgG antibody, HRP-conjugate (Sigma-Aldrich, 12-348), and goat-anti-mouse IgG antibody, HRP-conjugate (Thermo Fisher Scientific, 62-6520).

### RNA-seq on sorted cancer cells

Total RNA was prepared from FACS-sorted neoplastic cells ranging from 5 × 10^4^ to 4 × 10^5^ cells. RNA quality was assessed using the RNA 6000 Pico Assay Kit on the Agilent 2100 Bioanalyzer (Agilent). RNA used for RNA-seq had a mean RIN of 6.6. RNA-seq libraries were generated by Novogene, Inc. Total RNA (from 2.5-10 ng/sample) was used for cDNA synthesis using the Ovation RNA-seq System (NuGEN Technologies, Inc.). Two micrograms of NuGEN-amplified double-stranded cDNA was sheared using a Covaris sonicator to an average length of 490 bp and subjected to library preparation using the Takara V4 Ultra low input mRNA non-directional library preparation according to the manufacturer’s protocol and sequenced on a NovaSeq PE150 (∼6G raw data per sample).

### Analysis of RNA-seq data sets

Paired-end RNA-seq reads were aligned to the mm10 mouse genome using STAR under standard input parameters. Two *KT;Stag2^flox^* samples were excluded from subsequent analysis based on two parameters: 1) >20% the average number of nucleotide counts in Exon 8 of STAG2, and 2) >40% mRNA expression of *Stag2* from RNA-seq results (TPM). The differentially expressed genes between different tumor genotypes were called by DESeq2 using the HTSeq-derived counts as input (71). Unsupervised hierarchical clustering and heatmap visualization of DEGs were performed using the ‘pheatmap’ package in R (v.1.0.12, (1)).

The DESeq2-calculated fold changes were used to generate ranked gene lists for input into GSEA (v.4.0.3). GSEA was performed using normalized RNA-seq counts against gene signatures from the MSigDB database (72). Default parameters were used with the following exception: max size= 20,000. Plots were made using the Rtoolbox package in R (https://github.com/PeeperLab/Rtoolbox). clusterProfiler (v.4.3.1.900) was used to perform GO analyses, GO term enrichment, and functional enrichment analysis (73). GO terms were then categorized using EMBL’s European Bioinformatics Institute (EMBL-EBI) goslim_mouse annotations and statistics analysis by “all” relationships (“occurs_in”, “has_input”, etc.).

### ssGSEA Analysis of previously published data set

*KT* and *KT;p53^flox;flox^(KPT)* bulk RNA-seq samples were analyzed at various stages of tumor progression (55 samples) (44). Single sample GSEA was performed on all samples using the GSVA package v1.46.0 in R (74), and linear regression and unpaired t-tests were performed in GraphPad Prism.

### ATAC-seq library preparation and data analysis

After cells were sorted and counted, 50-100,000 cells were resuspended in 1 mL of cold ATAC-seq resuspension buffer (RSB; 10mM Tris-HCl pH 7.4, 10 mM NaCl, and 3 mM MgCl_2_ in water) (75). Cells were centrifuged 500 x g for 5 min at 4 °C. After centrifugation, 900 μL of supernatant was aspirated. The remaining 100 μL of supernatant was carefully aspirated with a P20 pipette tip to avoid disturbing the cell pellet. Cell pellets were then resuspended in 50 μL of ATAC-seq RSB containing 0.1% NP40, 0.1% Tween-20, and 0.01% digitonin and mixed by pipetting up and down five times. This reaction was incubated on ice for 3 minutes, and after lysis, 1 mL of ATAC-seq RSB containing 0.1% Tween-20 was added, and tubes were inverted six times to mix. Nuclei were then centrifuged for 10 min at 500 x g at 4 °C. Supernatant was again removed using the two-pipette steps, as described before, and nuclei were resuspended in 50 μL of transposition mix (25 μL 2X TD buffer, 2.5 μL transposase (100 nM final), 16.5 μL PBS, 0.5 μL 10% Tween-20, and 5 μL water) by pipetting up and down six times. Transposition reactions were incubated at 37 °C for 30 minutes at 1000 rpm. Reactions were purified using Qiagen MinElute Reaction Cleanup Kit. ATAC-seq library preparation was performed as described using the primers listed in Supplementary Table 10 and sequenced on the NovaSeq 6000 platform (Illumina) with 2 × 75 bp reads. Adapter-trimmed reads were aligned to the mm10 genome using Bowtie2 (2.1.0). Aligned reads were filtered for quality using SAMtools (v.1.9), duplicate fragments were removed using Picard (v.2.21.9-SNAPSHOT) and peaks were called using MACS2 (v.2.1.0.20150731) with a q-value cut-off of 0.01 and with a no-shift model. Peaks from replicates were merged, read counts were obtained using bedtools (v.2.17.0) and normalized using DESeq2 (v.1.26.0). For the *KT* versus *KT;Stag2^flox^* samples all libraries had a TSS enrichment greater than 10. For the *KT;H11^LSL-Cas9^* sgInert and sgPaxip1 samples all libraries had a TSS enrichment greater than 8.

### In situ Hi-C library construction on sorted cancer cells

*In situ* Hi-C libraries were prepared from FACS-sorted neoplastic cells with low input ranging from 2.5 × 10^5^ to 3.5 × 10^5^ cells. In detail, cells were first pelleted at 300 G and resuspended in 1 mL Room Temp PBS containing 3% BSA, then fixed as previously described, (76) only with a higher centrifuge speed at 2,500 G for pelleting the fixed cells. Fixed cell pellets were processed through the standard Arima Hi-C kit (Catalog # A510008), until the step of Covaris DNA sonication to an average size of 400 bp. Sheared DNA was purified and size-selected by a two-step AMPure XP bead (Beckman Coulter #A63882) cleanup at 0.6x and 1.0x, and then inputted into the 2S Plus DNA Library Kit from Integrated DNA Technologies (Catalog #10009878) for low-input library preparation, until the step before library PCR amplification. Libraries underwent a biotin pulldown using streptavidin to enrich chromatin interactions before an initial five-cycle amplification was performed and a qPCR quantification determining the number of additional cycles required for each library to reach a final concentration around 20 nM in 20 µL. Amplified libraries were then examined by Agilent 4200 TapeStation for confirmation of size distribution before submission for sequencing. Libraries were first sequenced using the Illumina MiniSeq PE50 for quality control (analysis described in the next section), and then deeply sequenced using the Illumina NovaSeq PE50 aiming at >3 × 10^8^ read pairs for each library that passed the MiniSeq QC.

### Analysis of in situ Hi-C data

Hi-C data was aligned to the mm10 reference genome using MWA-MEM (77). Reads were filtered (MAPQ >= 30) and paired using the previously described pipeline (78). PCR duplicate reads were detected and removed by Picard (https://broadinstitute.github.io/picard/). For MiniSeq quality control analysis, we sorted unique read pairs into three categories: 1) cis-reads that aligned to the same chromosome with a distance over 1 kb, 2) cis-reads that aligned to the same chromosome with a distance equal or less than 1 kb, and 3) trans-reads that aligned to two different chromosomes. Hi-C libraries with a ratio of cis-reads with distances over 1 kb larger than 50% and trans-reads ratio less than 25% were proceeded with deep sequencing using the NovaSeq.

For deeply sequenced Hi-C data, .hic files were generated and normalized (-w 5000) using JuiceBox (79) after alignment and duplication removal described above. Hi-C data from replicates within the same treatment were merged before visualization and loop-calling using JuiceBox. For loop-calling, genome-wide KR normalization and three resolutions were used (-k GW_KR -r 5000, 10000, 25000). Called loop lists from wildtype and *Stag2*-deficient neoplastic cells were compared to each other and sorted into indicated categories: 1) common loops that both loop anchors have overlap between *KT* and *KT;Stag2^flox^* samples, 2) unique loops that neither loop anchors have overlap between *KT* and *KT;Stag2^flox^* samples, 3) loops that are unique to *KT* that have one loop anchor that overlaps with a loop unique to *KT;Stag2^flox^*samples, 5) loops that are unique to *KT;Stag2^flox^* that have one loop anchor that overlaps with common loops between *KT* and *KT;Stag2^flox^* samples, 6) loops that are unique to *KT;Stag2^flox^* that have one loop anchor that over laps with a loop unique to *KT* samples (**Fig. 5e)**. For loop intensity analysis, loops from both *KT* and *KT;Stag2^flox^* samples were concatenated and removed for redundancy, before using the juicer dump function to output the ratio of observed/expected for every loop in each separate sample (**Supplementary Table 11**).

### Histology and IHC

Lung lobes were preserved in 4% formalin and embedded in paraffin. Hematoxylin and eosin stains were performed by Stanford Pathology/Histology Service Center (Stanford, CA) or Histo-Tec Laboratories (Hayward, CA). Total tumor burden (tumor area/total area × 100%) and individual tumor sizes were determined using ImageJ. Immunohistochemistry (IHC) was conducted on 4 μm sections. Antigen retrieval was performed with 10mM citrate buffer in a pressurized decloaking chamber. The slides were washed with 1X PBST and prepared with VECTASTAIN ABC-HRP Kit (Vector Laboratories, PK-4000). The following primary antibody and dilution was used: NKX2-1 (1:250, Abcam, ab7013). Sections were developed with DAB (Vector Laboratories, SK-4100) and counterstained with hematoxylin. To assess NKX2-1 expression in tumors while taking into account potential differences in staining between samples and across sections, we compared NKX2-1 staining intensity between tumors and adjacent normal tissue. Tumors were binned as having 1) higher expression relative to adjacent normal tissue, 2) equal expression relative to adjacent normal tissue, or 3) lower expression relative to adjacent normal tissue.

## Data Availability Statement

All data generated or analyzed during this study are included in this published article (and its supplementary information files). RNA-seq, ATAC-seq, and Hi-C metadata and raw sequencing files can also be found in the Gene Expression Omnibus (accession IDs: GSE274204, GSE274205, and X).

## Supplementary Material

**Supplementary Figure 1.**
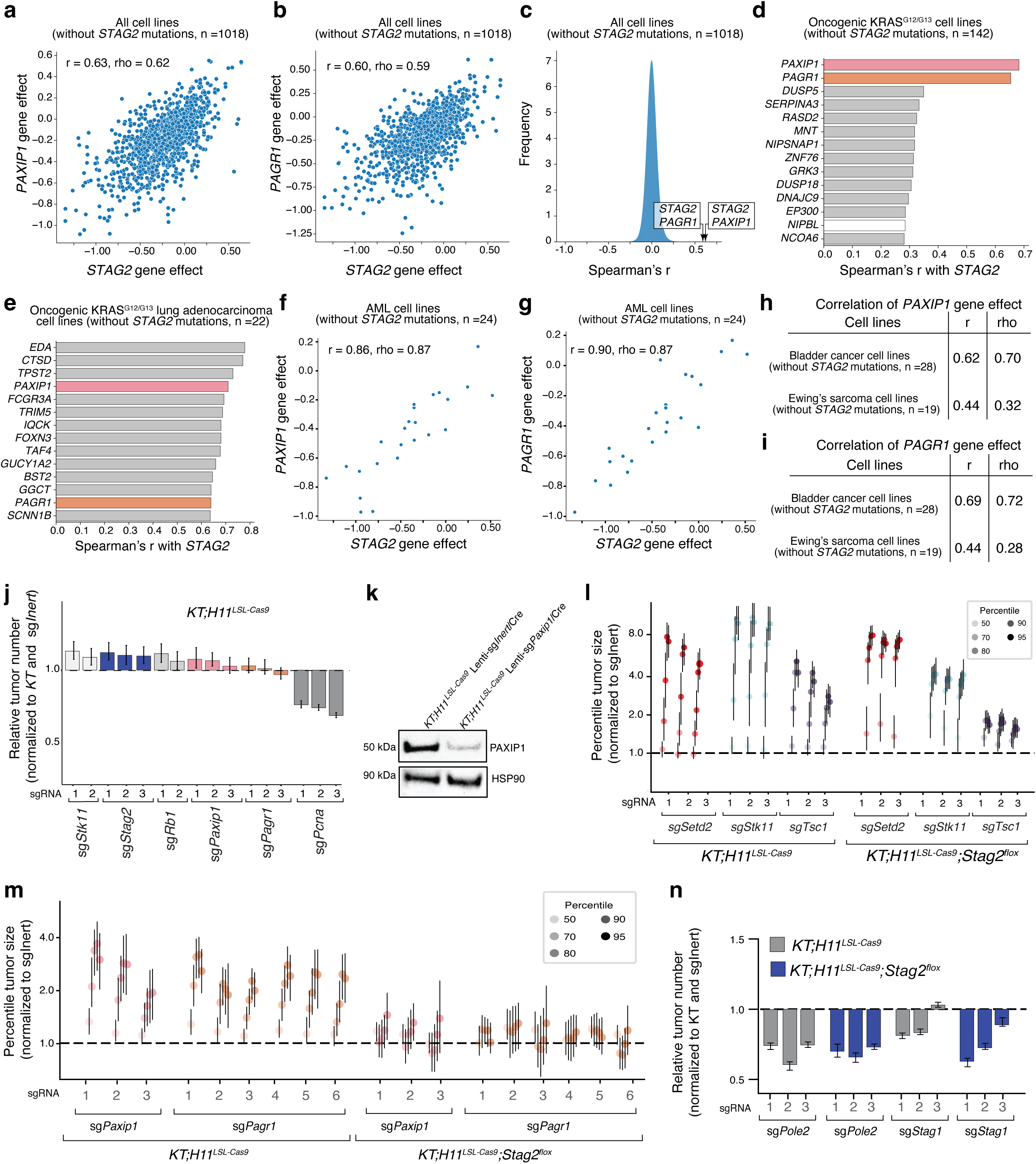
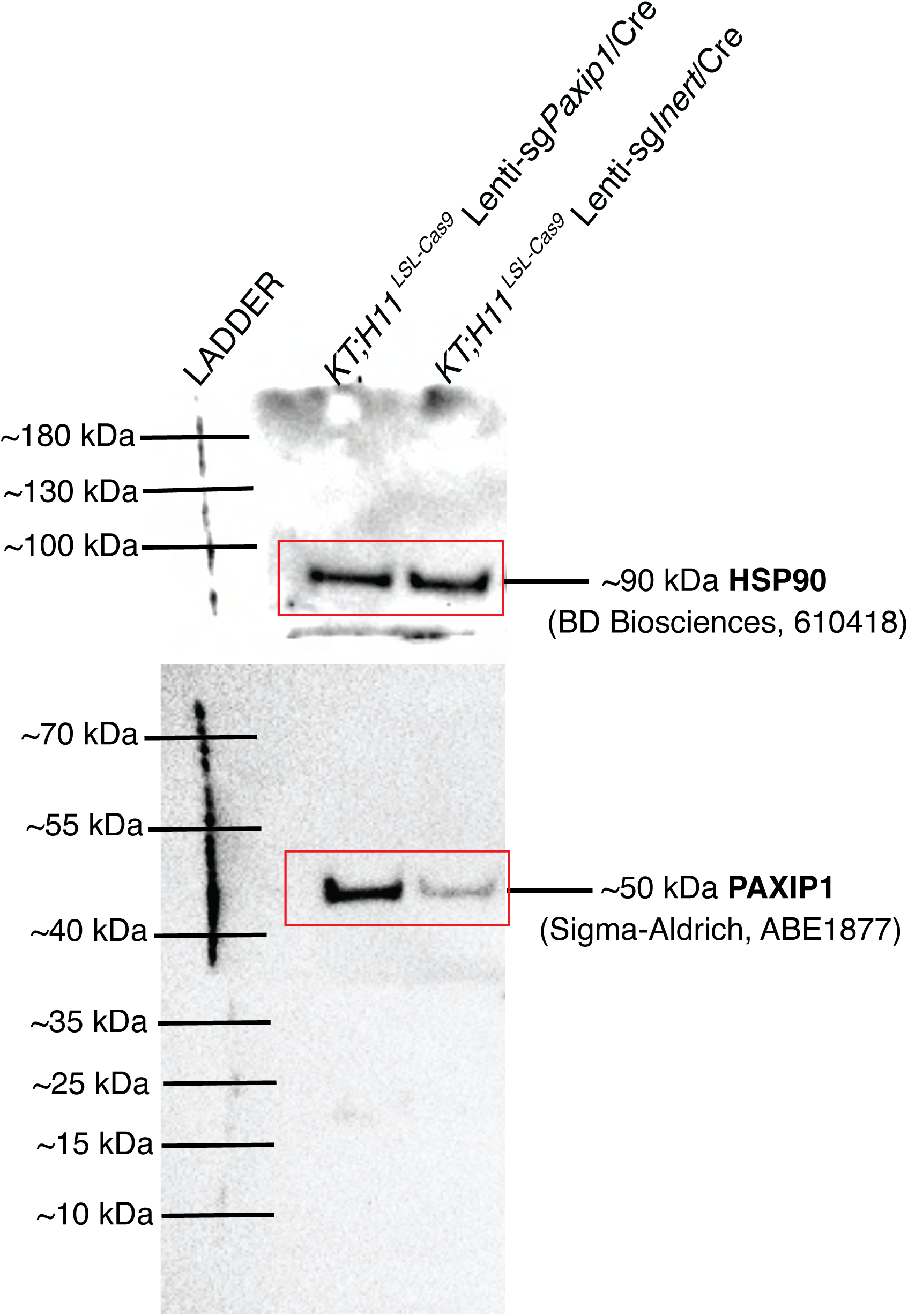
The effect of *STAG2* inactivation is highly correlated with that of *PAXIP1* and *PAGR1* inactivation in human cancer cell lines and genetic interactions with STAG2 are robust to different measures of tumorigenesis and tumor growth in oncogenic Kras lung cancer *in vivo*. **a-b)** Gene knockout effects for *PAXIP1* and *STAG2* inactivation (**a**) and for *PAGR1* and *STAG2* inactivation (**b**). Each dot represents a cell line. Cell lines with *STAG2* mutation were excluded. Spearman’s r and Pearson rho are indicated. **c)** Bell curve showing the frequency of Spearman’s correlations across all pairwise comparisons in DepMap. *STAG2-PAGR1* and *STAG2-PAXIP1* correlations are indicated. **d-e)** Genes with the highest Spearman’s correlation with the effect of *STAG2* inactivation from DepMap. Data from cell lines with oncogenic mutations at codons 12 or 13 of KRAS (**d**) and lung adenocarcinoma cell lines with oncogenic mutations at codons 12 or 13 of KRAS (**e**). Cell lines with *STAG2* mutation were excluded. *PAXIP1* and *PAGR1* are colored bars. Core cohesin complex genes are white bars. **f-g)** Gene effects for *PAXIP1* and *STAG2* inactivation (**f**) and for *PAGR1* and *STAG2* inactivation (**g**) in Acute Myeloid Leukemia (AML). Each dot represents a cell line. Cell lines with *STAG2* mutations were excluded. Spearman’s r and Pearson rho are indicated. **h-i)** Tables indicating Spearman’s r and Pearson rho for *PAXIP1* gene effect (**h**) and for *PAGR1* gene effect (**i**) in Bladder cancer and Ewing’s sarcoma. Cell lines with *STAG2* mutations were excluded. **j)** Relative tumor number (normalized to *KT* and sgInert). Mean +/− 95% confidence intervals are shown. Dotted line indicates no effect. Raw values and significance of each effect is shown in Supplemenary Table 2 (**j**). **k)** Western blot on sorted neoplastic cells from *KT;H11^LSL-Cas9^* mice with tumors initiated with the indicated Lenti-sgRNA/Cre vectors. Data represents one replicate of three independent experiments. **l)** Tumor sizes at the indicated percentiles for tumors with sgRNA targeting *Setd2, Stk11,* or *Tsc1* (normalized to sg*Inert*) in *KT;H11^LSL-Cas9^* and *KT;H11^LSL-Cas9^;Stag2^flox^* mice. Each gene was targeted with three sgRNAs. Error bars indicate 95% confidence intervals. Dotted line indicates no effect. **m)** Tumor sizes at the indicated percentiles for tumors with sgRNA targeting *Paxip1* or *Pagr1* (normalized to sg*Inert*) in *KT;H11^LSL-Cas9^* and *KT;H11^LSL-Cas9^;Stag2^flox^* mice. Each gene was targeted with three and six sgRNAs, respectively. Error bars indicate 95% confidence intervals. Dotted line indicates no effect. **n)** Comparison of relative tumor number for tumors with sgRNAs targeting for *Stag1, Stag2,* or *Pole2* in *KT;H11^LSL-Cas9^* mice compared to *KT;H11^LSL-Cas9^;Stag2^flox^* mice. Mean +/− 95% confidence intervals are shown. Raw values and significance of each effect is shown in Supplementary Table 4 (**l-n**).

**Supplementary Figure 2.**
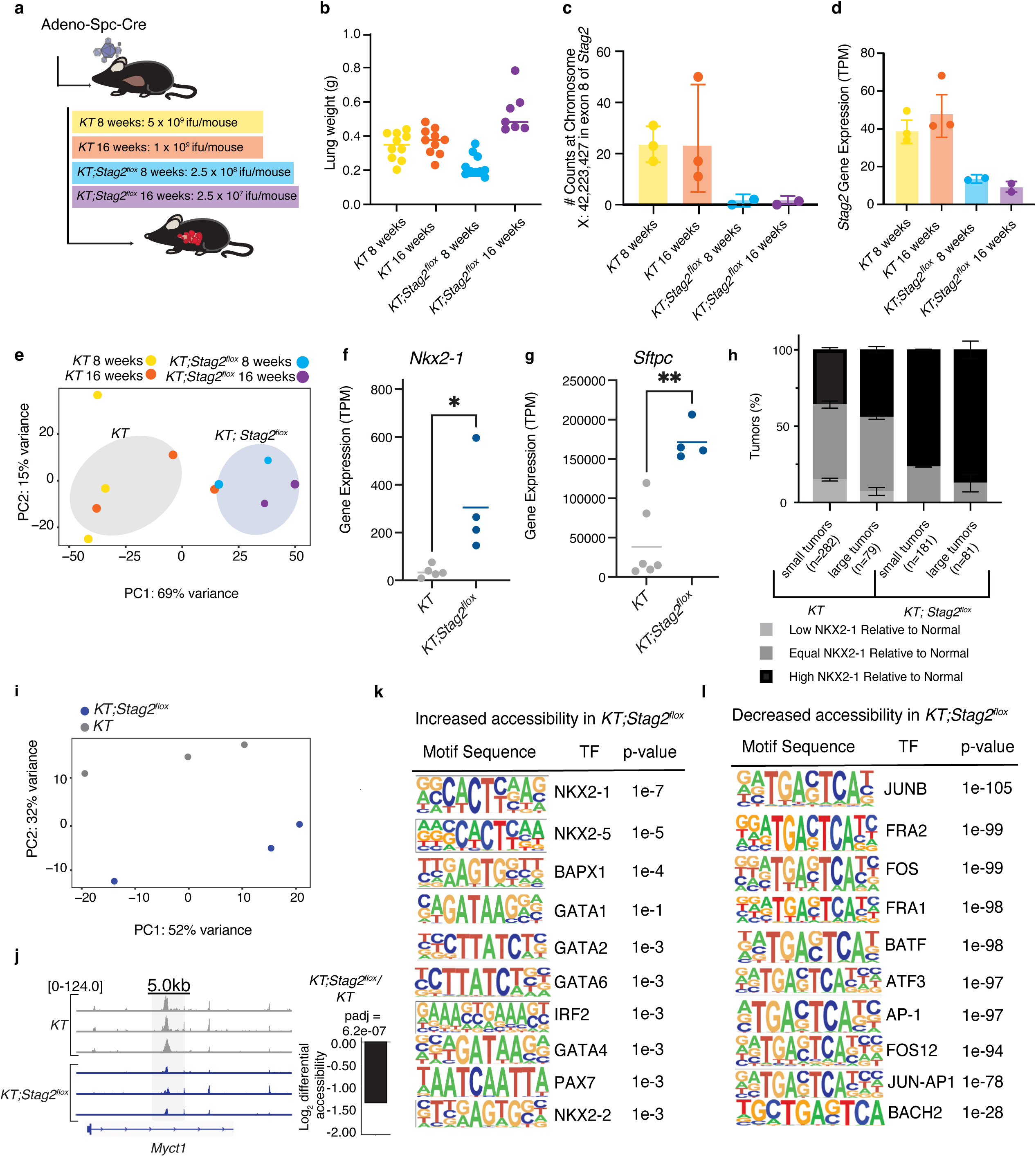
Validation of STAG2 inactivation, clustering of *KT* and *KT;Stag2^flox^* samples, gene expression of surfactant genes, and differential accessibility regulated by STAG2. **a)** Schematic of tumor initiation with Adeno-Spc-Cre with different viral titers in *KT* and *KT;Stag2^flox^* mice to generate tumor samples to analyze at 8 weeks and 16 weeks after tumor initiation. **b)** Lung weights of mice in each group. Each dot represents a mouse and the bar is the mean. **c)** Number of DNA nucleotide counts at Chromosome X locus in Exon 8 of STAG2 in each group. Note that this region is within floxed exon. Mean +/− SEM is shown. **d)** *Stag2* RNA expression in TPM via RNA-seq in each group. Bar is the standard error. Mean +/− SEM is shown. **e)** Principal component analysis (PCA) of *KT;Stag2^flox^* and *KT* tumors processed with RNA-seq libraries at 8 weeks and 16 weeks clustered by genotype. **f-g)** Gene expression (TPM) for *Nkx2-1* (**f**) and *Sftpc* (**g**) for *KT* and *KT;Stag2^flox^* 8 weeks and 16 weeks. ** p-value < 0.01, *\** p-value < 0.1 by unpaired t-test. Each dot represents a mouse and the bar is the mean. **h)** Quantification of NKX2-1 expression in *KT;Stag2^flox^* (n=2 mice) and *KT* (n=2 mice) tumors at 8 weeks after initiation.Tumor NKX2-1 expression quantified by comparison to NKX2-1 expression in adjacent normal tissue. Tumor number quantified labeled in graph. Mean +/− SD is shown (p-value <0.001 by Ordinary one-way ANOVA) **i)** Principal component analysis (PCA) of *KT* and *KT;Stag2^flox^* tumors processed with ATAC-seq libraries at 16 weeks clustered by genotype. ATAC-seq signal tracks for the *Myct1* gene locus in tumors from *KT* and *KT;Stag2^flox^* mice. **j)** Motif sequences and corresponding p-values for transcription factors with motif enrichment in regions with increased accessibility in cancer cells from *KT;Stag2^flox^* mice. **k)** Motif sequences and corresponding p-values for transcription factors with motif enrichment in regions with decreased accessibility in cancer cells from *KT;Stag2^flox^* mice. **l)** Motif sequences and corresponding p-values for transcription factors with motif enrichment in regions with decreased accessibility in cancer cells from *KT;Stag2^flox^* mice.

**Supplementary Figure 3.**
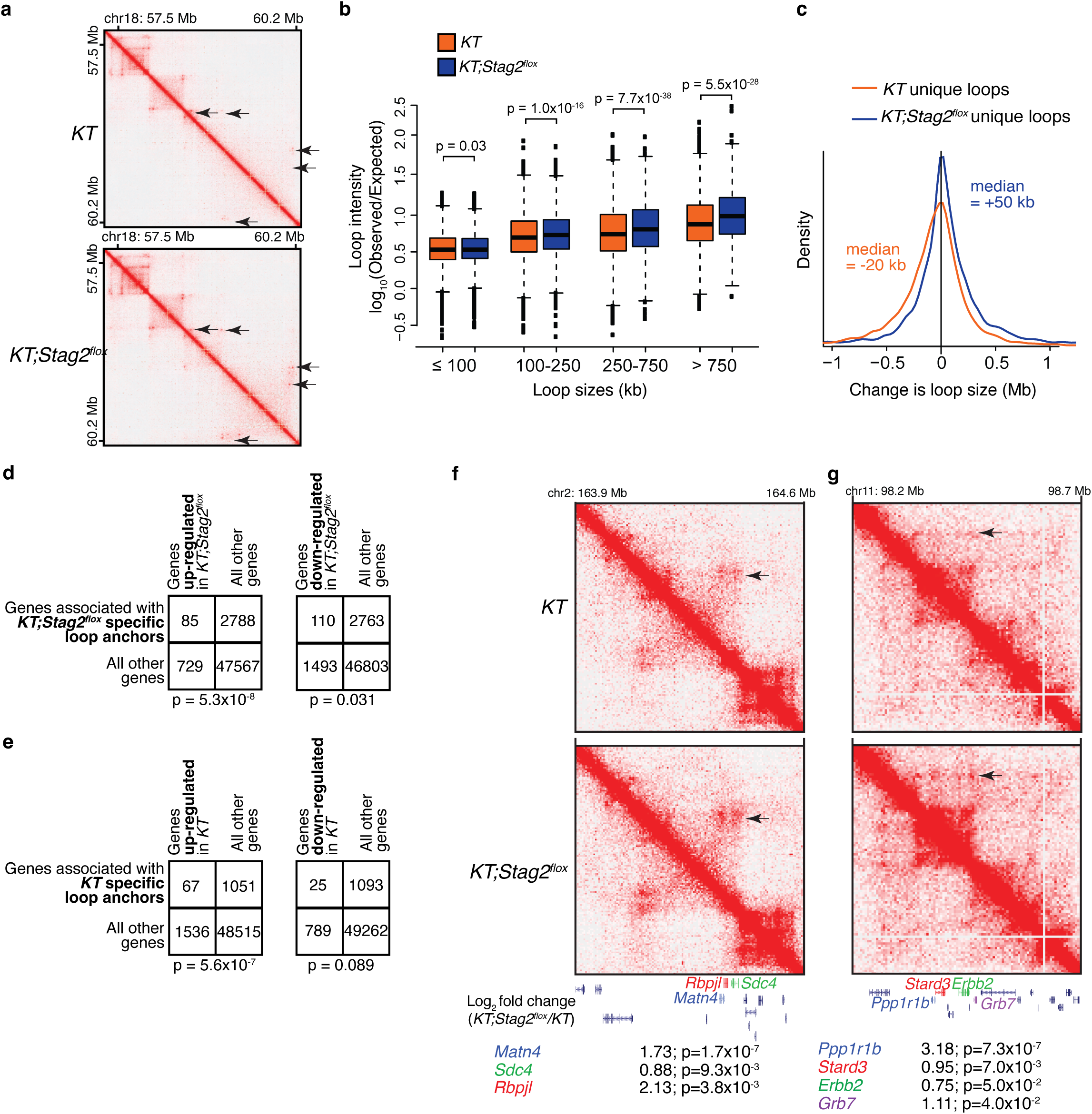
*KT* and *KT;Stag2^flox^* samples contain both unique and common DNA loops and DNA anchors that correlate with gene expression but not chromatin accessibility. **a)** Normalized contact matrices in *KT* (top) and *KT;Stag2^flox^* (bottom) neoplastic cells. Contact matrices were visualized in Juicebox with the same color scale. Arrows indicate novel or stronger loop contacts in *KT;Stag2^flox^*. **b)** Comparison of loop intensity (log_10_ (Observed/Expected)) of the indicated loop sizes between all loops in *KT* and *KT;Stag2^flox^* samples. Boxes show median +/− interquartile range. Whiskers show standard error. P-values (Wilcoxon rank test) are shown. **c)** Change in loop size between the *KT;Stag2^flox^* unqiue loops and the common loops or *KT* unqiue loops with which they share one anchor (blue). Change in loop size between *KT* unique loops and the common loops or *KT;Stag2^flox^* unqiue loops with which they share one anchor (orange). **d)** Number of genes associated with *KT;Stag2^flox^* unique loops that are up-regulated (left) or down-regulated (right) in *KT;Stag2^flox^* samples versus all other genes that are and are not associated with *KT;Stag2^flox^* unique loops. p-value (chi-square test) is shown. **e)** Number of genes associated with *KT* unique loops that are up-regulated (left) or down-regulated (right) in *KT* samples versus all other genes that are and are not associated with *KT* unique loops. p-value (chi-square test) is shown. **f,g)** Differences in chromatin looping and gene expression (from RNA-seq data, log_2_ fold change (*KT;Stag2^flox^ / KT*); p-value) of several differentially expressed genes promixal to unique anchor sites.

**Supplementary Figure 4.**
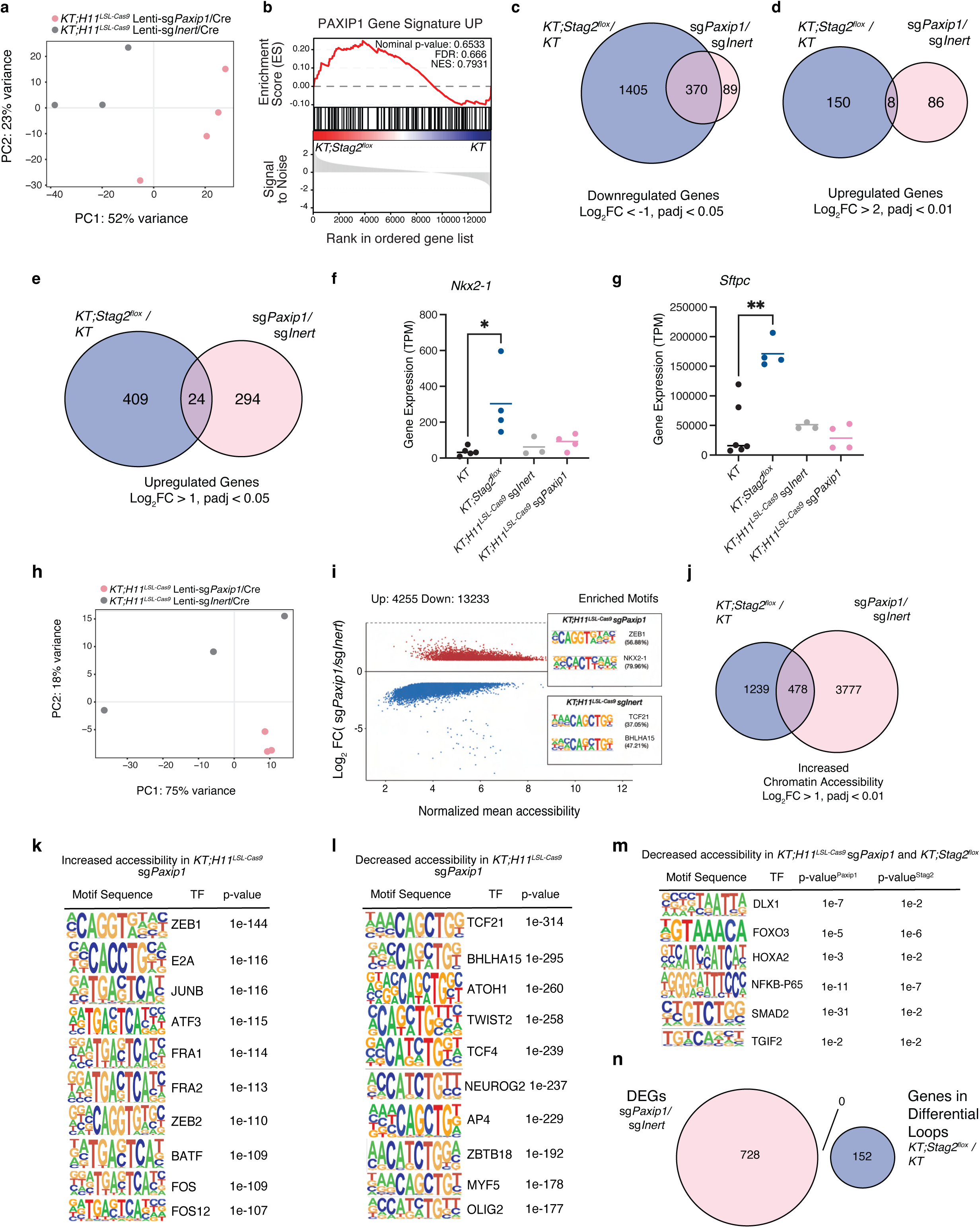
PAXIP1 and STAG2-cohesin regulation of gene expression and chromatin accessibility is conserved in downregulation but not upregulation of genes. **a)** Principal component analysis (PCA) of RNA-seq on *KT;H11^LSL-Cas9^* sg*Inert* and *KT;H11^LSL-Cas9^* sg*Paxip1* tumors. **b)** Upregulated PAXIP1 gene signature enrichment in rank-ordered gene list for *KT* versus *KT;Stag2^flox^*. **c-e)** Venn diagram of shared downregulated genes (log_2_FC < −1, padj < 0.05) (**c**), shared upregulated genes (log_2_FC > 2, padj < 0.01) (**d**), and shared upregulated genes (log_2_ FC > 1, padj < 0.05) (**e**) between *KT;Stag2^flox^/KT* and *KT;H11^LSL-Cas9^* sg*Paxip1*/*KT;H11^LSL-Cas9^* sg*Inert*. **f-g)** Gene expression (TPM) for *Nkx2-1* (**f**) and *Sftpc* (**g**) in neoplastic cells from tumors from *KT, KT;Stag2^flox^, KT;H11^LSL-Cas9^* sg*Inert,* and *KT;H11^LSL-Cas9^*sg*Paxip1* in mice. Each dot is an RNA-seq sample and the bar is the mean. ** p-value < 0.01, *\** p-value < 0.1 by unpaired t-test (*KT* and *KT;Stag2^flox^* data are same as Supplementary Fig. S2). **h)** Principal component analysis (PCA) of *KT;H11^LSL-Cas9^* sg*Inert* and *KT;H11^LSL-Cas9^* sg*Paxip1* tumors processed with ATAC-seq libraries. **i)** Differential accessibility across 17488 significant peaks in *KT;H11^LSL-Cas9^ sgInert* and *KT;H11^LSL-Cas9^* sg*Paxip1* mice (3 mice/group). The x-axis represents the log_2_mean accessibility per peak and the y-axis represents the log_2_FC in accessibility. Colored dots are significant (log_2_FC > |1|,FDR < 0.05). Red dots are increased chromatin accessibility in *KT;H11^LSL-Cas9^* sg*Paxip1* accompanied by transcription factor hypergeometric motif enrichment in *KT;H11^LSL-Cas9^* sg*Paxip1*, and blue dots are decreased chromatin accessibility in *KT;H11^LSL-Cas9^* sg*Paxip1* accompanied by transcription factor hypergeometric motif enrichment in *KT;H11^LSL-Cas9^* sg*Inert*. **j)** Venn diagram of shared regions of increased chromatin accessibility (log_2_FC > 1, padj < 0.01) between *KT;Stag2^flox^/KT* and *KT;H11^LSL-Cas9^* sg*Paxip1*/ *KT;H11^LSL-Cas9^* sg*Inert*. **k-l)** Motif sequences and corresponding p-values for transcription factors with motif enrichment in regions with increased accessibility in *KT;H11^LSL-Cas9^* sg*Paxip1* (**k**) and with decreased accessibility in *KT;H11^LSL-Cas9^* sg*Paxip1* (**l**). **l)** Motif sequences and corresponding p-values for transcription factors with motif enrichment in regions with decreases accessibility in *KT;H11^LSL-Cas9^* sg*Paxip1* and *KT;Stag2^flox^*. **m)** Venn diagram of differentially expressed genes regulated by *KT;H11^LSL-Cas9^* sg*Paxip1*/*KT;H11^LSL-Cas9^* sg*Inert* (|log_2_ FC| > 1, padj < 0.05; RNA-seq) versus genes found in differential loops regulated by *KT;Stag2^flox^/KT* (HiC).

**Supplementary Figure 5.**
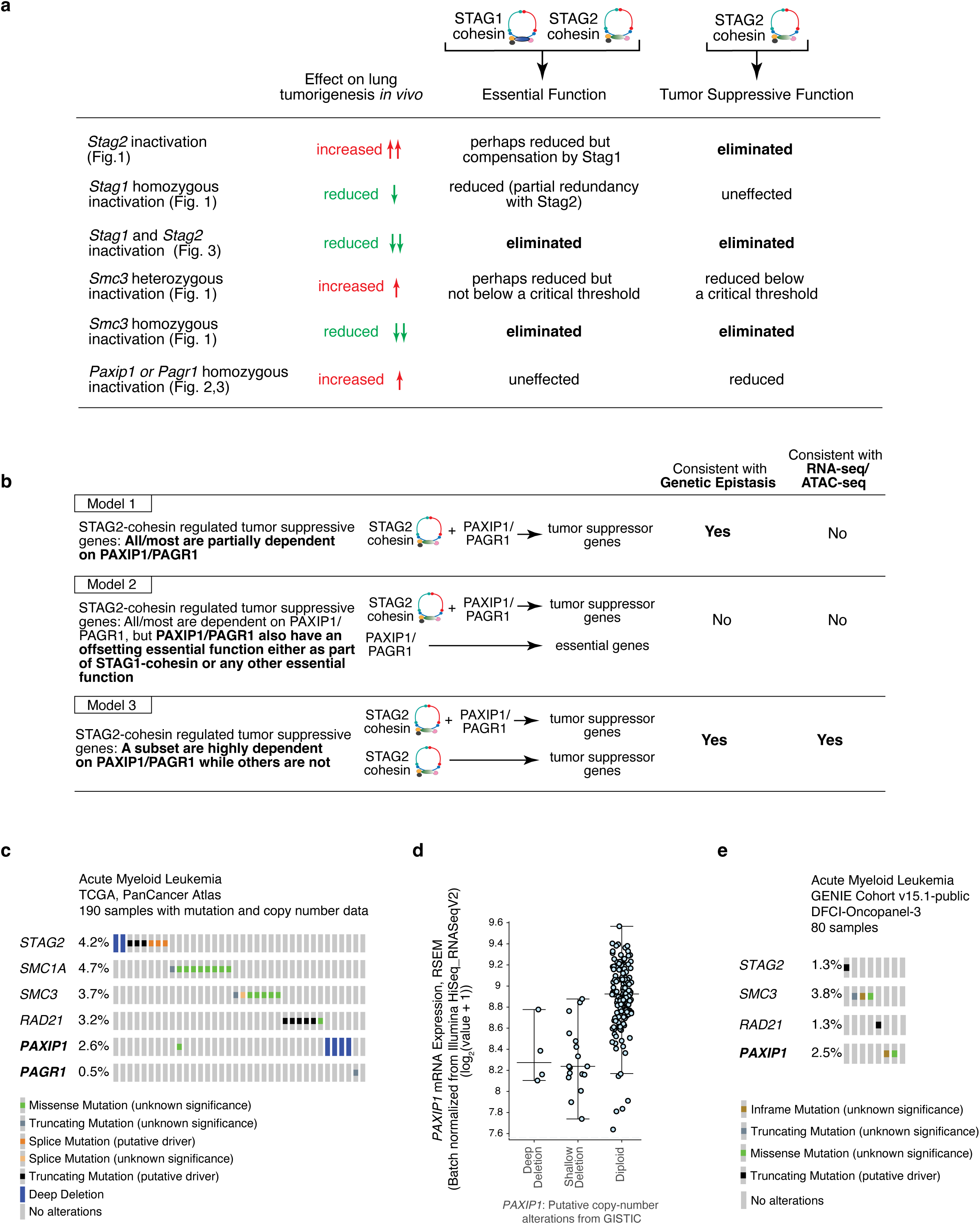
Summary of the *in vivo* genetic experiments as they related to the essential role of cohesin in general and the specific tumor suppressive function of STAG2-cohesin and PAXIP1/PAGR1, as well as a summary of the most parsimonious model of STAG2-cohesin and PAXIP1/PAGR1 mediated tumor suppression and mutations in *PAXIP1* and *PAGR1* and downregulation of *PAXIP1* mRNA expression in a subset of acute myeloid leukemias. **a)** Interpretations of results from Figures 1-3 describing the distinct phenotypes controlled by STAG2- and STAG1-cohesin, homozygous and heterozygous inactivation of cohesin components, and the role of PAXIP1 on lung tumorigenesis *in vivo.* The lung tumor suppressive effect of Stag2 is well established in oncogenic KRAS-driven lung tumors (Cai *et al.*, 2021 and Blair *et al.*, 2023), confirmed in the current study, and extended to lung cancer driven by other oncogenes (Blair *et al.*, 2023). The compensation of STAG1 and STAG2 in the essential functions of cohesin, in general, is well established in cell lines (Arruda *et al.*, 2020, van der Lelij *et al.*, 2017, Canudas & Smith *et al.*, 2009) and confirmed in lung cancer by our *in vivo* studies, including the genetic epistasis between *Stag1* and *Stag2*. While homozygous inactivation of each core and auxiliary cohesin component greatly reduced lung tumorigenesis, heterozygous inactivation of *Smc3* using a floxed allele increases tumorigenesis. This likely explains the mutations in cohesin components in human cancer and extends the importance of dysregulation of this complex to a much larger fraction of lung adenocarcinomas. Finally, while *Paxip1* or *Pagr1* inactivation increased lung tumorigenesis, inactivation of either gene did not reduce tumorigenesis of Stag2-deficient tumors, as would have been expected if *Paxip1/Pagr1* were also involved in the essential cohesin function. **b)** Multiple models could have explained how STAG2-cohesin and the PAXIP1/PAGR1 complex cooperate to suppress lung tumorigenesis. However, our genetic epistasis data (Figure 3) and molecular analyses (Figures 4-5) are most consistent with Model 3 in which the major role of PAXIP1/PAGR1 is to work with STAG2-cohesin to regulate a subset of genes that are controlled by STAG2-cohesin. **c)** Oncoprint of acute myeloid leukemias from TCGA accessed through cBioPortal. 190 samples with mutation and copy number data. Mutation type is indicated. **d)** mRNA expression of *PAXIP1* in acute myeloid leukemias from TCGA accessed through cBioPortal. 165 samples with mutation, copy number, and gene expression data. Sample were split based on putative copy number of *PAXIP1*. Each dot is a sample. Note low expression in a subset of sample with likely unaltered DNA copy number (diploid samples). **e)** Oncoprint of acute myeloid leukemias from DCFI-Oncopanel-3 samples from GENIE accessed through cBioPortal. 80 samples with mutation data. *SMC1A* and *PAGR1* were not profiled. Mutation type is indicated.

**Table S1** shows tumor size and number metrics from the cohesin complex member Tuba-seq^Ultra^ analysis. **Table S2** shows tumor size and number metrics from *Paxip1, Pagr1, Cdca5*, and various TSGs and essential genes Tuba-seq^Ultra^ analysis. **Table S3** shows genes Selected for Tuba-seq^Ultra^ analysis of putative genetic interactors of STAG2. **Table S4** shows tumor size and number metrics from Tuba-seq^Ultra^ analysis of putative genetic interactors of STAG2. **Table S5** shows metrics and RLE-normalized (DESeq2) counts for *KT;Stag2^flox^* versus *KT* 8 weeks and 16 weeks after tumor initiation RNA-seq experiment. **Table S6** shows Metrics and RLE-normalized (DESeq2) counts for *KT;H11^LSL-Cas9^* sg*Paxip1* vs *KT;H11^LSL-Cas9^* sg*Inert* RNA-seq experiment. **Table S7** shows metrics for differentially expressed genes (DEGs) altered by both STAG2 and PAXIP1 with Log_2_FoldChange > (abs) 1. **Table S8** shows metrics and RLE-normalized (DESeq2) counts for *KT;Stag2^flox^* versus *KT* 16 weeks after tumor initiation ATAC-seq experiment. **Table S9** shows metrics and RLE-normalized (DESeq2) counts for *KT;H11^LSL-Cas9^* sg*Paxip1* versus *KT;H11^LSL-Cas9^* sg*Inert* 16 weeks after tumor initiation ATAC-seq experiment. **Table S10** shows a list of forward and reverse primers used to generate ATACseq libraries for *KT;Stag2^flox^* versus *KT* and KT;H11^LSL-Cas9^ sg*Paxip1* versus KT;H11^LSL-Cas9^ sg*Inert* ATACseq experiments. **Table S11** shows loci for all loops and normalized metrics for *KT;Stag2^flox^*versus *KT* 16 weeks after tumor initiation Hi-C experiment.

